# Automated Analysis of Bird Head Motion from Videos in Unconstrained Settings

**DOI:** 10.1101/2024.12.20.629664

**Authors:** Marco Santos-Lopes, Ricardo Araújo, Romain David, Paulo L. Correia

## Abstract

This study introduces a framework applicable to videos of habitual behaviors of multiple bird species to automatically assess head angular velocities and frequencies during various behaviors in natural habitats. The process involves detecting birds, identifying key points on their heads, and tracking changes in their positions over time. Bird detection and key point extraction were trained on publicly available datasets, including Animal Kingdom, NABirds, Birdsnap, CUB-200-2011, and eBird, featuring videos and images of diverse bird species in uncontrolled settings. Initial challenges arose due to the complexity of video backgrounds, leading to misidentifications and inaccurate key point estimates. These issues were addressed through validation, refinement, filtering, and smoothing steps. Head angular velocities and rotation frequencies were computed from the refined key points. The algorithm performed well at moderate speeds but was limited by the 30 Hz frame rate of most eBird videos, which constrained measurable angular velocities and frequencies and caused motion blur, affecting key point detection. Our findings suggest that the framework may provide plausible estimates of head motion but also emphasize the importance of high frame rate videos in future research, including extensive comparisons against ground truth data, to fully characterize bird head movements.

## 1. Introduction

As agile organisms, birds are expected to present fast head motion, detected by their semicircular duct system. This sensory organ, enclosed inside bony semicircular canals and located in the vestibular system of the inner ear, monitors head rotations and is essential for vision, navigation, spatial awareness, balance and motor coordination [1-6]. Semicircular canals have often been used to predict locomotion, behavior, ecology and, more recently, body temperature in fossil vertebrates [1-11]. However, to date, the extreme scarcity in kinematic data still hampers our understanding of the actual relationship between semicircular duct function and head motion.

In this context, this study aims to lay the groundwork for automatically analyzing bird head motion from videos. By doing so, it will provide the foundational information necessary for future studies to gather sufficiently accurate data to better understand how bird head motion relates to the functional limits of their semicircular duct system [1], with an approach that could be extended to other vertebrate species in the future.

This approach belongs to the field of pose estimation, where Bird Pose Estimation (BPE) plays a pivotal role. BPE is a computer vision task focused on detecting and analyzing the orientation and position of birds or their body parts within an image or video [12]. This process provides spatial information, typically represented by key points, or landmarks, in 2D or 3D. Previous studies have applied BPE to birds [13, 14] using identified key points to classify their behavior. In this work, the contribution lies in applying 2D BPE to a bird’s head, hereby defined as Bird Head Pose Estimation (BHPE).

Deep neural networks (DNNs) [15–17], such as HRNet, have been effectively used to enhance the accuracy and applicability of human pose estimation [18]. Hence, in this work we also used the HRNet architecture for BHPE, which required the availability of an extensive training dataset covering a wide range of bird species, morphologies, poses, backgrounds, and lighting conditions. However, the development of bird-specific datasets is still in its infancy, and most available datasets provide a limited number of images with 2D annotated key points. This scarcity of data can lead to considerable prediction errors in DNN outputs [19, 20] and constrains BHPE to the 2D realm for now. To start addressing this issue, this work contributes a newly developed manually annotated dataset based on eBird videos [21], enabling the calculation of head motion parameters, such as angular velocity and frequency, for a large range of bird species, along with the proposal of a methodology to automatically detect birds in unconstrained scenarios, and estimate, refine and validate a set of 2D key points to describe the bird’s head motion, with applicability to the 3D realm.

Additionally, the DNNs used for key point estimation require a fixed-size input, preferably with the bird centered in the video frames. To achieve this, an object detection module was included for bird detection, and the selection has been to use the well-known YOLO [23], as it is very effective in object detection problems, providing bounding boxes (BBs) and locating them in the image. However, the limited availability of bird datasets can result in object detection errors when considering videos captured in unconstrained settings, as well as the diversified morphologies of birds, which this work minimizes by proposing a detection validation module.

In summary, the main contributions of this work are the:

- Proposal of a bird detection module, based on the YOLO model, specifically trained to identify birds in video footage acquired in unconstrained settings, followed by a validation module, specifically developed to improve detection results and their stability along time;
- Development of a HRNet model, which was initially proposed for human pose estimation, to be applied to the problem of 2D bird head pose estimation in videos captured in unconstrained scenarios;
- Proposal of a methodology for computing bird head motion metrics from videos, including the identification of the minimal set of bird head key points required for this purpose;
- Making available to the research community: a comprehensive dataset, entitled BirdGaze, tailored for BHPE models, using online sourced datasets and manually annotated frames from eBird videos, and including metadata such as the BB center coordinates, a scale factor to apply for resizing the BB, and the bird head key point locations.

## 2. Materials and Methods

This project proposes a new framework, depicted in Figure 1, for the automatic computation of birds’ head angular velocities and frequencies, using videos as the source for head motion.

**Figure 1.**
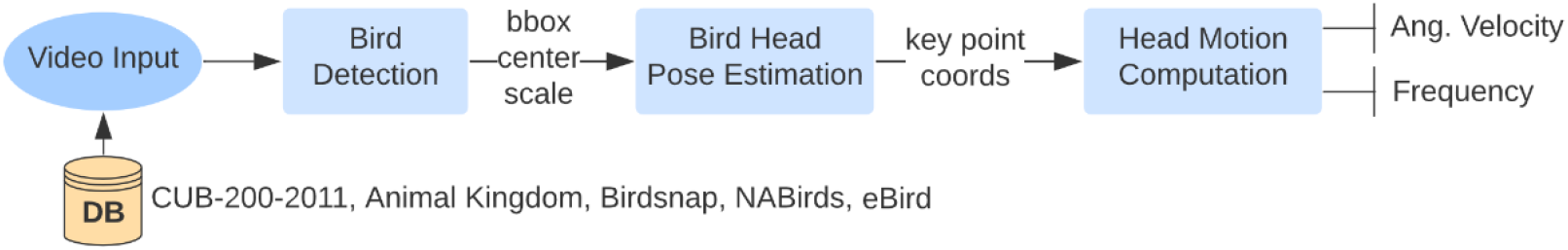
Architecture of the proposed bird head motion video analysis system.

It consists of three main modules, namely:

- **A Bird Detection Module**: In this first module, the video is processed as a set of frames. Bird detection is carried out using the YOLO model [24], trained and validated on CUB-200-2011 [25] and tested on Birdsnap [26] datasets. Bounding boxes are positioned on each frame, around detected birds, and validated using a new trajectory filter we developed to improve results. Each frame containing an isolated bird is then cropped along the bounding box, centered and scaled.
- **A Bird Head Pose Estimation Module**: In this second module, scaled frames are inputted into the BHPE model, a modified version of the HRNet model [18], which provides 2D coordinates of birds’ head key points. This model has been trained on the Animal Kingdom [27], NABirds [28], Birdsnap [26] and eBird [21] datasets, and tested on eBird images derived from videos not included in the training set. Key points are validated using a set of methods we specifically developed to handle extreme bird morphologies (e.g., toucans or owls), to minimize erroneous key point estimations, and to reduce instabilities in noisy key point positions, ensuring improved and smoother estimations.
- **A Head Motion Computation Module**: In this last module, validated birds’ head key points are used to compute head angular velocity at each frame, as the angular change in head orientation between two consecutive frames, divided by elapsed time. The module also describes the head motion frequency spectrum as the power spectral density (PSD) calculated via a Fourier analysis of the magnitude and direction of head rotation through time.

### 2.1. Bird Datasets

Annotated bird data is essential for training and evaluating bird detection and BHPE models, which will allow extracting head motion data for the birds observed in the videos. Although bird-specific datasets are still in the preliminary stages of development, several relevant publicly available datasets have been identified and used in this work, as listed in Table 1.

**Table 1.** Summary of used bird datasets.

| Datasets | Training<br>(images) | Validation<br>(images) | Testing<br>(images) | Annotations | Task |
| --- | --- | --- | --- | --- | --- |
| eBird [21] * | 3,147 | - | 107<br>images | Center<br>Scale<br>2D Key Points | BHPE<br>Bird Detection<br>Bird Head Motion |
| Animal Kingdom<br>[27] | 6,819 | 1,705 | - | Center<br>Scale<br>2D Key Points | BHPE |
| CUB200-2011 [25] | 5,993 | 2,897 | - | BB | Bird Detection |
| NABirds [28] | 48,000 | - | - | Center<br>Scale<br>2D Key Points | BHPE |
| Birdsnap [26] | 47,386 | - | 1,704 | Center<br>Scale<br>2D Key Points<br>BB | BHPE<br>Bird Detection |

\* All testing images were extracted from eBird videos not included in the training stage.

For the development of the bird detection module, the dataset considered comprises 5,993 training images from CUB200-2011 [25], 2,897 validation images from CUB200-2011 and 1,704 validation images from Birdsnap [26], all of which contain annotations for the position of bounding boxes around the birds present in the images. CUB200-2011 was selected for its diverse content in terms of bird species (200) and large image count, while Birdsnap provides samples for distinct bird species, to evaluate model generalization. The training and validation splits follow the original partitions of the CUB-200-2011 dataset [25], while the testing partition is based on the original test set of the Birdsnap dataset [26]. These original splits were considered appropriate for model training and preserving samples for generalization validation.

For the development of the bird head pose estimation module a new 2D BHPE annotated dataset is proposed, here entitled **BirdGaze**, which includes images from four prominent sources: the Animal Kingdom [27], NABirds [28], Birdsnap [26] and eBird [21]. These datasets represent the largest publicly available collections and are widely recognized in the literature for their significant role in avian research [34,35]. Their extensive morphological diversity is crucial for this study. Besides the bird images, the proposed BirdGaze dataset includes a set of annotations, notably:

- Center of the bounding box containing the bird body, described by its 2D coordinates;
- Scale factor, defining a multiplying factor to apply to the bird bounding box for resizing it to fit a fixed rectangle size, which is used as input to the adopted key point extraction model;
- Coordinates of the four selected 2D key points: top of head, tip of beak, left eye, right eye.

As part of this work, it was necessary to identify the minimum number of key points needed for angular velocity and frequency computations. After several experimentations, and checking which key points could be automatically identified with reliability for a large number of bird species with different morphologies, the decision was to select four key points: the top of the head (i.e., crown), left eye, right eye, and beak (i.e., beak tip) – using these points it is possible to determine a bird’s head gaze direction. Figure 2 illustrates an example from the BirdGaze dataset, highlighting the key points annotated on the bird’s head.

**Figure 2.**
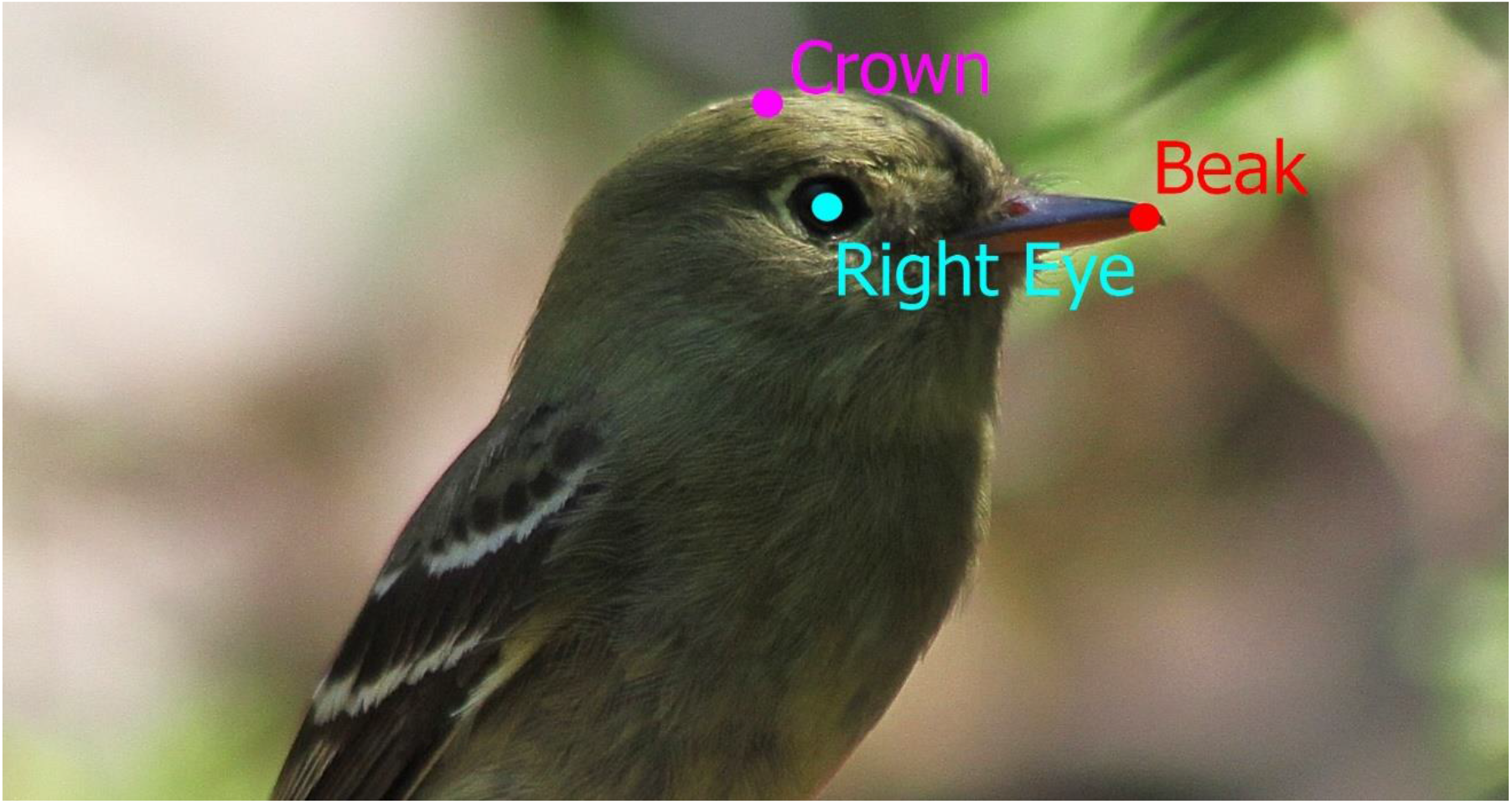
Example from the BirdGaze dataset, showcasing annotations.

In addition to the mentioned data, the BirdGaze dataset also includes 3,254 manually annotated frames extracted from eBird videos [21]. From these manually annotated frames, 3,147 images were used for training fine-tuning, extracted from the following videos: 8 videos of *Anas undulata*, 4 of *Apus apus*, 9 of *Asio otus*, 7 of *Caprimulgus europaeus*, 8 of *Ciconia ciconia*, 8 of *Meleagris gallopavo*, 8 of *Phoenicopterus ruber*, 9 of *Pteroglossus bailloni*, 8 of *Spheniscus demersus*, 8 of *Struthio camelus*, and 10 of *Tyto alba*. These species were selected because they are morphologically diverse, and their membranous labyrinth is known [1]. The 107 testing images were randomly selected from different videos of the 72 bird species in the eBird video dataset. The final BirdGaze dataset thus consists of a total of 104,705 images allocated for training, 1,705 for validation, and 1,811 for testing. This substantial training set is essential for maximizing the accuracy of estimated bird head key points, thereby enhancing the precision of the subsequent bird head motion computations.

### 2.2. Bird Detection

Figure 3 shows the main processing steps of the bird detection module.

**Figure 3.**
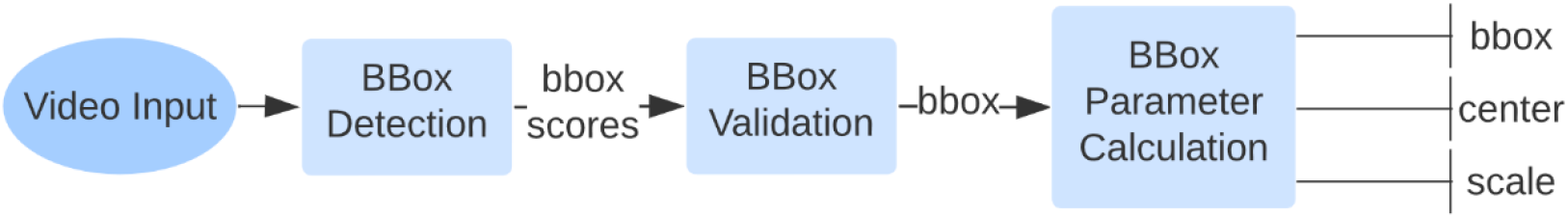
Main processing steps of the Bird Detection module.

The YOLO architecture is widely recognized as one of the best object detection models available [23, 29, 30]. This led to the adoption of YOLOv8 [24], which, after training, demonstrated high accuracy and fast execution. For each bird detected in a video frame YOLOv8 outputs an associated BB, as well as a confidence score.

A BB validation method is then applied to verify that the detected BBs correctly represent birds. The proposed validation method relies on a filter that checks the consistency of a bird’s trajectory throughout the video, checking whether a detected BB follows a continuous path along time. This validation filter includes three main steps:

- **Confidence score thresholding**: Detected BBs with a YOLOv8 confidence score below 80% are removed, to retain only high-confidence bird detections. This results in a mix of genuine detections, missed detections, and hopefully a very limited number of false positive detections.
- **Trajectory validation and completion**: It is expected that genuine detections form continuous and smooth sequences along the video timeline. A set of continuous sequence candidates to represent each bird’s trajectory is identified, using the Intersection over Union (IoU) metric to assess the similarity between BBs in consecutive time instants. Trajectories corresponding to genuine bird detections, are selected from the set of candidate sequences by comparing with the original YOLOv8 detected BBs, also considering those with lower detection confidence scores. BBs with high IoU values are incorporated into the trajectory, including for video frames where detections were previously missing. BBs that were not initially detected by YOLOv8 but are necessary to maintain a consistent temporal sequence are added through interpolation.

After obtaining the BB around the bird for each video frame, the center and a scale factor are computed, to normalize the input image for the subsequent BHPE module.

- The center is calculated as the midpoint of the bird’s bounding box.
- The scale factor is computed considering the BB size and the default input size expected by the key point estimation model adopted.

An affine transformation is then applied by the BPHE module, which uses the center and scale to crop and resize the bird’s BB to match the fixed input size expected by the key point estimation model.

### 2.3. Bird Head Pose Estimation

The architecture of the BHPE module is shown in Figure 4. Each of the main processing steps is detailed below.

**Figure 4.**
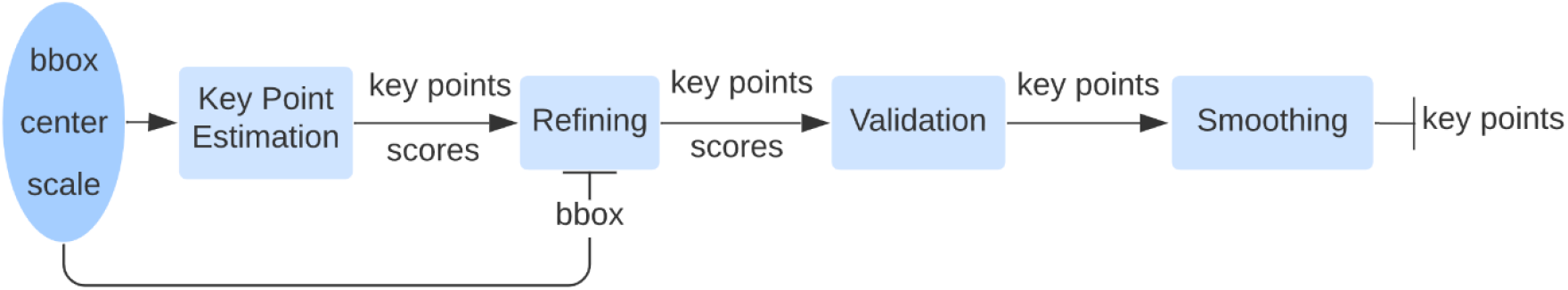
Main processing steps of the BHPE module.

#### 2.3.1. Key Point Estimation

Bird head pose estimation starts with the 2D key point estimation step, which takes as input a centered, scaled and cropped region of the original video frame, using the parameters computed by the bird detection module. This ensures that the bird images input to the BHPE model have the expected dimensions. The BHPE model then estimates the four selected key points: top of the head, left eye, right eye, and beak, along with a set of associated confidence scores.

Initial experiments with BHPE architectures, such as ResNet-50 [31], resulted in high key point location error rates, leading to the adoption of a more complex but more powerful architecture, the HRNet [19]. The first model developed using this architecture, named **HRNet4**, corresponds to a modified version of the original HRNet to identify the desired four bird head key points. HRNet4 was trained using the Animal Kingdom dataset and, although it resulted in a better performance than when using ResNet50, it was still lacking in the ability to compute reliably the four key points. This prompted the expansion of the dataset, and the creation of the proposed BirdGaze dataset. Training HRNet4 with this expanded dataset, lead to a new model, named HRNetExp. While HRNetExp showed promising results, it still struggled with occlusions and complex image backgrounds. Further attempts were then made using a more complex architecture, the VHRNet [33], an enhanced version of HRNet, employing transformers to exploit the potential of self-attention mechanisms. However, the trained VHRNet model did not surpass the performance previously achieved with HRNetExp, due to constraints in the size of the training dataset set, and to limitations in the available computational infrastructure. Consequently, **HRNetExp** was chosen as the final BHPE model for this work.

To minimize errors in key point estimations, three additional steps were proposed, for key point refining, validation and smoothing. While these steps enhance key point accuracy and filter out errors, some inaccuracies remain due to the rapid head rotations that may lead to unusual bird poses and cause motion blur, which HRNetExp struggled to manage.

To address the unusual bird pose issue, further eBird videos showcasing diverse bird morphologies and head poses were manually annotated and added to the training set. The BirdGaze dataset, now incorporating eBird, Animal Kingdom, NABirds, and Birdsnap, was used to fine-tune HRNetExp, enhancing its ability to handle fast head rotations and unusual poses, thus improving bird head motion computation.

#### 2.3.2. Bounding Box and Key Point Refining

The BB and key point refining step addresses issues where certain bird morphologies, such as the elongated necks of ostriches, lead to the head being cropped out of the frame, resulting in incorrect estimations; this often occurs due to the identification of the bird’s body as the center, excluding the head from the cropped BB area.

To solve this, the proposed method adjusts the BBs to focus specifically on the bird’s head. The proposed procedure is illustrated in Figure 5.

**Figure 5.**
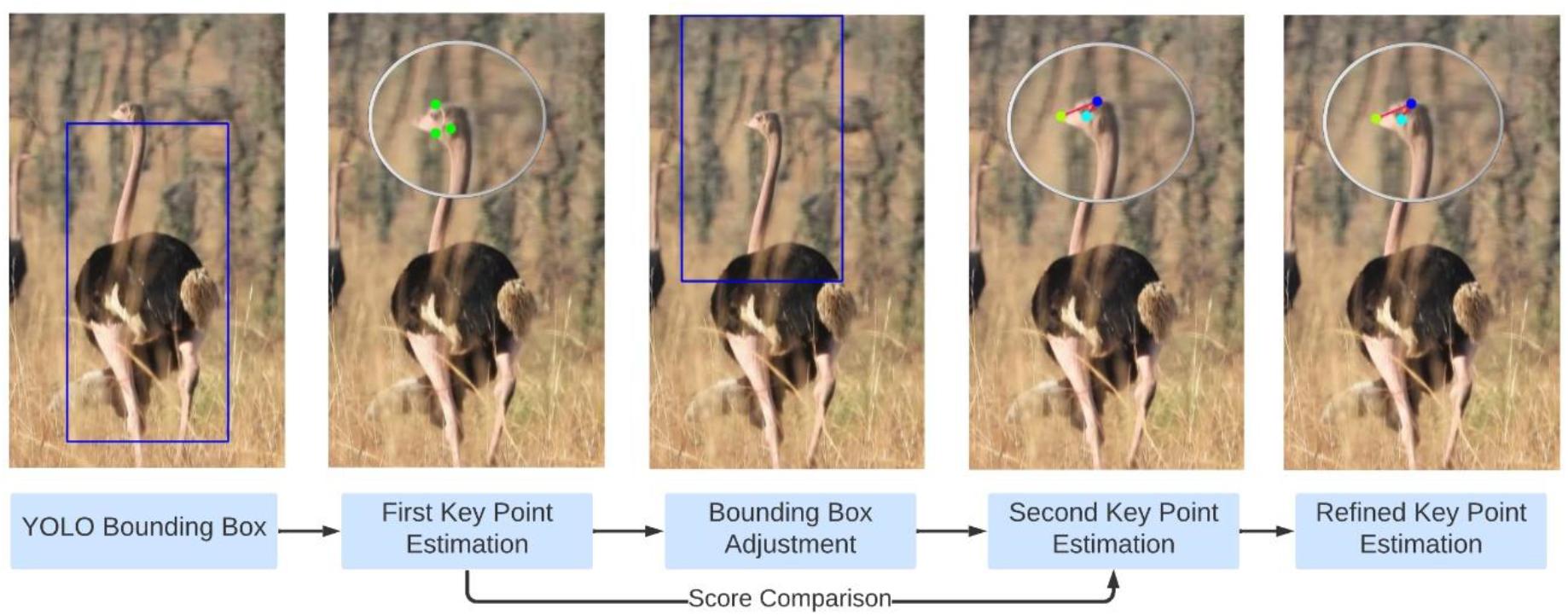
Illustration of the procedure proposed for BB and key point refining.

The proposed BB and key point refining procedure begins with the YOLOv8 bird detection that outputs body centered BBs, which allows a first iteration of key point estimation. Despite some incorrect estimations occurring due to the bird head not being included in the initial BB provided by the YOLOv8 model, the head centroid estimation computes a fair estimation of the head position. The bird detection BB is then adjusted to be centered on the head of the bird. Then, a second round of 2D key point estimation is conducted within these head-centered bounding boxes. Finally, these results are combined with those of the first round by selecting the highest confidence key point locations. This approach retains correct initial estimations while correcting errors due head exclusion in species with elongated necks.

#### 2.3.2. Key Point Validation

The key point validation step aims to identify and eliminate erroneous key point location estimations, which often arise when the model encounters unseen bird morphologies that were not present in the training set. Erroneous key point placements within the image are difficult to detect, as for the case of abrupt head movements, correctly estimated key points may resemble these inaccuracies. To mitigate the impact of such errors, a procedure is proposed that checks the following two constraints:

- **Sufficient HRNetExp confidence scores**: Key point estimations for which the HRNetExp confidence score is below 30% are discarded. This allows to filter out most of the erroneous key point estimations.
- **Consistency of key points with bird morphology**: Since the distance from the top of the head to each eye must not exceed the distance to the beak, to comply with all major bird morphologies, the proposed procedure removes key point estimations that violate this constraint.

While these procedures reduce errors, the first one is unable to deal with errors that are consistent along time. Therefore, an additional adaptive filtering mechanism is proposed, applying a maximum/minimum score threshold and a distance threshold.

This filter compares each estimation to the previous one:

- If the current estimation shows a large deviation from the previous one (based on the distance threshold) and its confidence score (from HRNetExp) is below a minimum threshold, it is removed, and a flag is set indicating possible consecutive errors in the upcoming estimations.
- In the next estimation, the filter checks the flag and whether that estimation has again a low HRNetExp confidence score. If so, this confirms a consecutive error, and the current estimation is removed.
- However, if the current estimation has a high HRNetExp confidence score (above the maximum score threshold) and deviates significantly from the previous estimation, this signals the end of consecutive errors. The current estimation is kept, and the flag is untoggled.

The maximum/minimum score and distance thresholds are determined using the same formula by varying an adjustment factor *p*. This approach leverages the cumulative moving average of confidence scores and key point distances across frames, and is updated as follows:

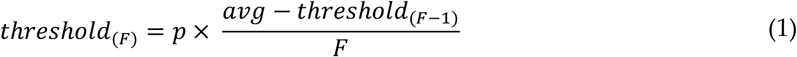

In this equation, *avg* represents the average distance or score and F represent the index of the frame. For distances, it is calculated by averaging the measured distances of the current estimated key points with those of the key points estimated in the previous time instant. For scores, the thresholds are computed by averaging the HRNetExp confidence scores of the current estimated key points. *threshold* denotes each of the computed average thresholds. *frame*_*index*_ corresponds to the specific frame number, and *p* is an adjustment factor that, when varied, allows to compute the maximum/minimum score thresholds and distance threshold, using the same equation. It is empirically set to 1.5 for the minimum score threshold, 2.4 for the maximum score threshold, and 0.05 for the distance threshold.

After calculating the above threshold values, they are passed through a sigmoid function to constrain the output value. The sigmoid function constrains the maximum/minimum score thresholds between 0 and 1, and the distance threshold between 0 and a value equal to *min*(*image*_*width*_, *image*_*height*_) pixels:

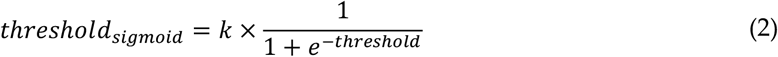

In this equation, *k* is the limiting value, set to 1 for scores and *min*(*image*_*width*_, *image*_*height*_) for distances. Regardless of the value of *p* in Equation 1, the *threshold*_*sigmoid*_ will always be constrained between 0 and *k*. A larger value of *p* will result in thresholds closer to 1 for scores and *min*(*image*_*width*_, *image*_*height*_) for distances.

This approach removes the need for hardcoded thresholds, allowing for dynamic adjustments based on factors like bird species, pose, and camera distance.

#### 2.3.3. Key Point Position Smoothing

This step aims to reduce flickering, which occurs when key points are slightly misaligned between consecutive frames. Flickering refers to inconsistent variations in the predicted positions of the key points across consecutive frames, despite the actual position of the key point remaining static or exhibiting minimal movement. This effect is caused by slight inaccuracies in the model’s prediction causing a flickering effect, thus introducing noise. This noise affects head motion calculations and is more noticeable in frequency calculations, especially for higher frequencies. This poses a problem, limiting the detection of head motion, because some frequencies cannot be reliably obtained because of noise. Although this noise cannot be fully removed, a filter can be applied to minimize its impact.

The proposed filter is based on angular displacement and key point proximity. It checks if the key point positions in the current frame deviate from those in the previous frame by calculating the angle between vectors formed by the head and beak key points and comparing those against an angle threshold. A threshold of 5° was empirically chosen to smooth out smaller and less significant head adjustments while maintaining responsiveness. However, it is important to note that this threshold’s effectiveness depends heavily on the video frame rate, as higher frame rates may inadvertently exclude actual head motions. A more robust approach might involve smoothing the power spectrum instead of the key points. Additionally, the maximum distance between consecutive homologous key points is measured to ensure it does not exceed a distance threshold based on the bird’s distance from the camera and bird morphology.

The distance threshold is calculated as follows:

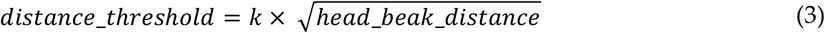

where *k* is a scalar value constrained by 0 < *k* ≤ 1. Increasing *k* allows for smoother outputs. A value of *k* = 0.4 was empirically found to work well across bird species with different morphologies. If both the angle and distance changes are within their thresholds, minimal movement is assumed, allowing for the smoothing of the key point position coordinates, defined by:

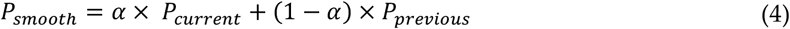

where *P* denotes the 2D coordinates of a key points. *α* is a smoothing factor, constrained by 0 < *α* ≤ 1, in which a smaller value gives more weight to the previous frame, creating a smoother effect. A larger value gives more weight to the current frame, making the output more responsive. A value of *α* = 0.15 was empirically found to provide better smoothing based on visual observation. This method reduced flickering, resulting in smoother head pose estimations.

### 2.4. Head Motion Computation

To analyze head motion, it is essential to set up a head centered frame of reference, characterized by fixed points on the head—such as the tip of the beak, the eyes, or the crown— and to track their change in position and orientation from a frame to the next. In this study, to compute angular velocity (i.e., the speed at which the head rotate), we first calculate the angular displacement θ as the scalar product between vectors linking key points of the crown and beak in successive frames:

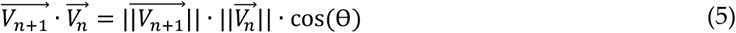

where *V*_*n*+1_ and *V*_*n*_ are the vectors for current and previous frames. Angular velocity *w* is then calculated as:

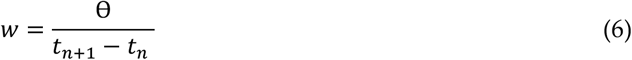

where *t*_*n*_ and *t*_*n*+1_ are the timestamps of consecutive frames.

Similarly, to compute head rotation frequency (i.e., how often the head rotate), we first calculate a scalar *R*, representing the magnitude and direction of head rotation, as the cross product between vectors linking key points of the crown and beak in successive frames:

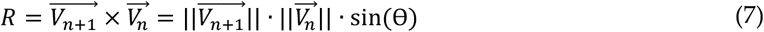

The power spectral density (PSD) of head rotation frequency *f* is then calculated from a time series of *R* as:

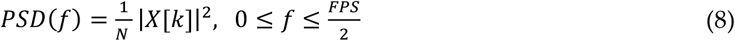

where *N* is the total number of frames of the video, *FPS* refers to the framerate of the video, and *X*[*k*] is the discrete Fourier transform (DFT) of the time series of *R*. According to the Nyquist theorem, measurable head rotation frequency cannot exceed half the framerate of analyzed videos.

The proposed methodology is primarily designed to analyze head motion using 2D key points from the head, which requires the viewing axis of input videos to be aligned with the pitch, roll or yaw axes of head rotation for results to be accurate. Most eBird videos capture birds in side view, hence limiting the analyses to the pitch rotations in this work. However, the methodology can be extended and validated in the 3D domain, allowing for the analysis of pitch, roll and yaw motion. Here we use the 3D-POP dataset [33], which contains videos of pigeons with annotated 3D key points, to validate the proposed methodology in the 3D domain. To compute pitch, roll, and yaw rotations in the 3D domain, we first define a reference frame on the pigeon head such that the vector *x* corresponds to the pitch axis of rotation of the head, and the vectors *y* and *z* to the yaw and roll axes of rotation, respectively. To do so we use the axis passing through the beak and nose key points as the roll axis and the one passing through the eye’s key points as the pitch axis. Both axes were normalized to obtained the unit pitch and roll vectors and the yaw vector was calculated by doing their cross-product. These vectors components are described within the world reference system such that for example 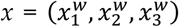. We then analyze how the orientation of the head reference frame changes between two consecutive video frames, *n* and *n* + 1. When transitioning from video frame *n* to frame *n* + 1, the change in head orientation is described by a transformation matrix. This matrix includes rotation and translation components. We isolate the rotation part of this matrix, which consists of three vectors 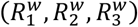, that define how the orientation changes within the world reference system. To compute a specific rotation (e.g., pitch), we perform a dot product between the rotation vector 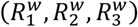 and the corresponding head rotation vector from video frame *n* in the world reference system. For pitch rotation:

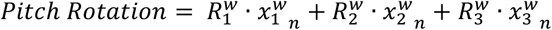

The same principle applies for yaw and roll, using their respective directional vectors in frame *n*. This validation enables a more comprehensive analysis, covering the full range of head motions.

## 3. Results

This section presents results for each module described in Figure 1, namely the bird detection module (section 3.1), the bird head pose estimation module (section 3.2), and the head motion computation module (section 3.3).

### 3.1. Bird Detection

In this section, the bird detection approach is presented, using the YOLOv8 model. Sub subsection 3.1.1 details the training, validation, and testing of YOLOv8 on the CUB-200-2011 dataset [25], focusing on model performance metrics such as Intersection over Union (IoU), precision, and recall. Sub subsection 3.1.2 covers the validation of the detected bounding boxes using eBird videos [21], highlighting the filtering technique used to smooth trajectories and correct errors caused by background complexity and occlusions.

#### 3.1.1. YOLOv8 training, validation and testing

Figure 6 illustrates the training progress of the YOLOv8 bird detection model over 1000 epochs using the CUB-200-2011 dataset [25], with accuracy assessed through the IoU metric, which measures the overlap between predicted bounding boxes and ground-truth annotations. The figure also displays “box_loss,” which indicates the accuracy of bounding box predictions, “precision,” which measures the proportion of correct bird detections among positive predictions, “recall,” which reflects the model’s ability to detect actual bird instances, and “mAP50,” which evaluates localization accuracy at a 50% IoU threshold.

**Figure 6.**
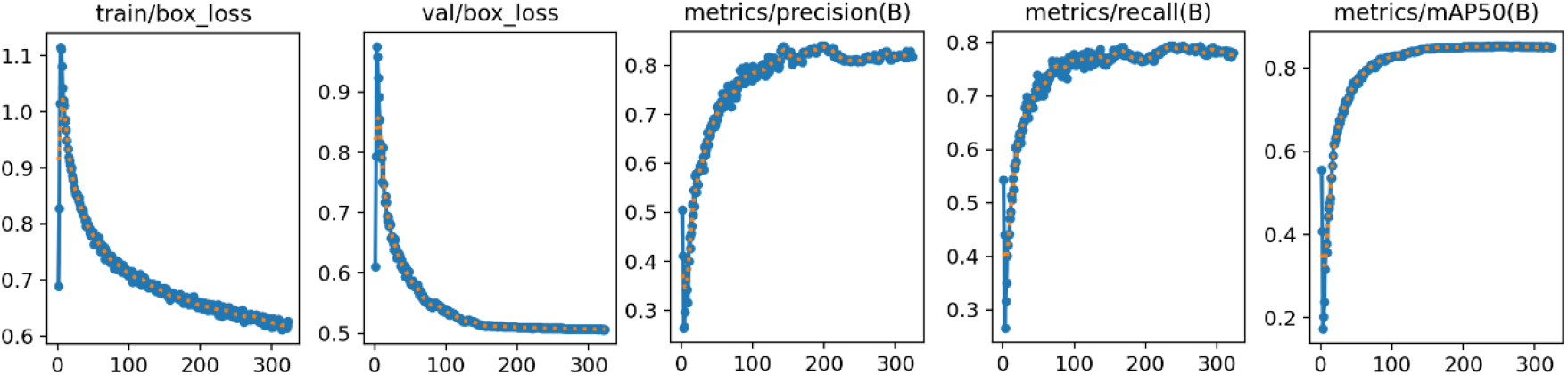
Training progress of YOLOv8, illustrating the evolution of training/validation loss, precision, and recall across epochs.

YOLOv8 demonstrated high precision and recall in bird detection, reaching a plateau after 323 epochs, indicating effective training on the CUB-200-2011 [25] dataset with 5,993 training images and 2,897 validation images, without any signs of overfitting. To evaluate the trained YOLOv8 model, the Birdsnap [26] test subset was used, and the average IoU results are summarized in Table 2.

**Table 2.** IoU scores obtained by trained YOLOv8 per image on Birdsnap [26] test subset.

| IoU Range | IoU > 0.75 | 0.75 ≥ IoU > 0.5 | 0.5 ≥ IoU > 0.25 | 0.25 ≥ IoU |
| --- | --- | --- | --- | --- |
| Number of Images | 1459 (86%) | 160 (9.43%) | 27 (1.59%) | 50 (2.95%) |

These results show that 86% of the YOLOv8 predictions on the Birdsnap test set achieved an IoU score of at least 0.75, indicating a good bird bounding box detection performance. A closer analysis of the images with low IoU scores (below 0.25) revealed that YOLOv8 had difficulties with occlusions and birds blending into their surrounding environment.

#### 3.1.2. Bounding Box Validation

Figure 7 illustrates the proposed bounding box validation procedure with an example trajectory of detected bounding boxes from an eBird video (ML599679051) featuring a long-eared owl (*Asio otus*) perched on a tree branch.

**Figure 7.**
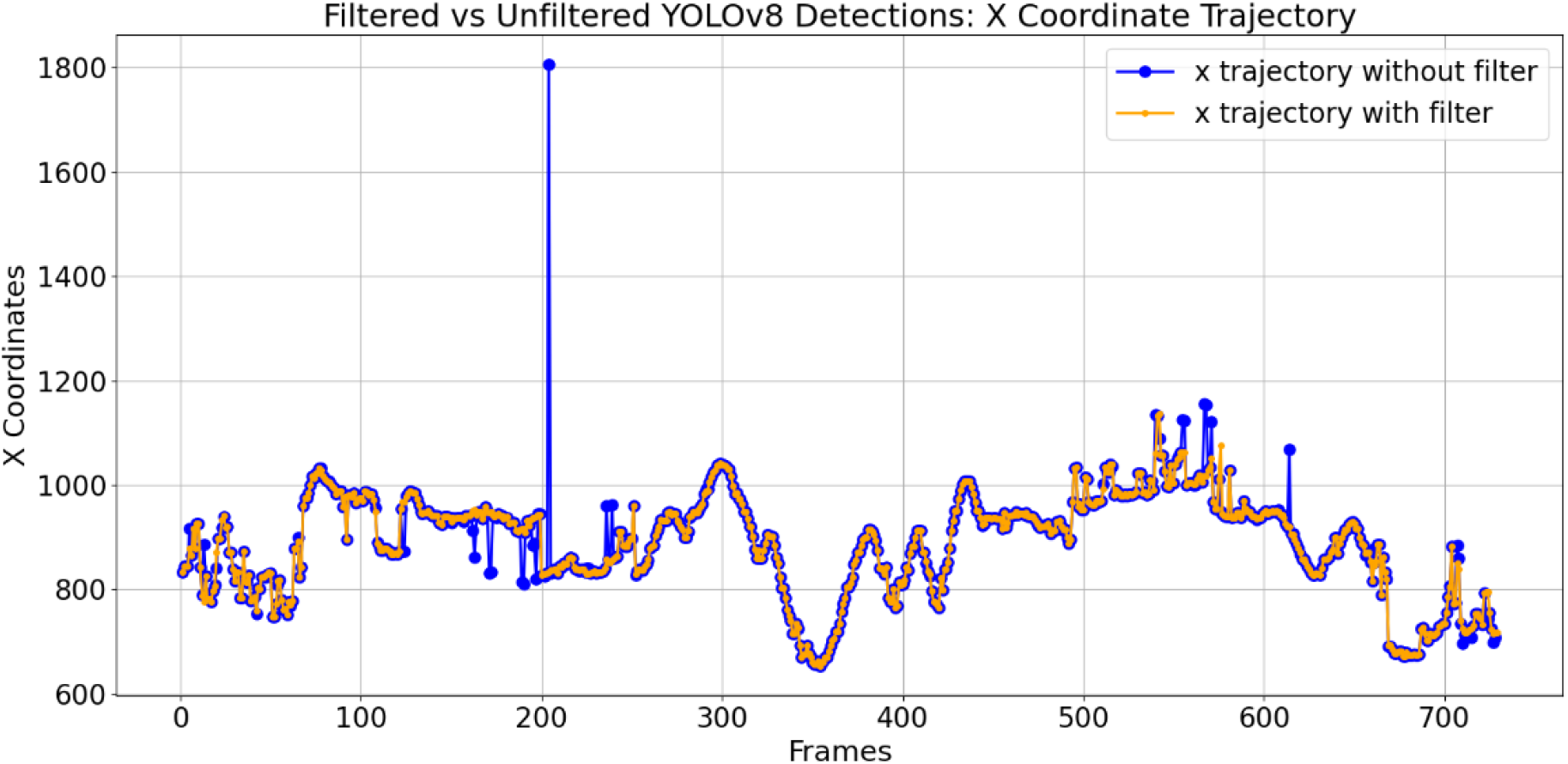
Trajectory of bounding boxes center along the X-axis across video frames for a long-eared owl.

Figure 7 displays the bird detection results, with the blue line representing unfiltered detections, including missing, correct, and incorrect estimated bounding boxes, where absent points and spikes indicate errors caused by background complexity. The yellow line represents the filtered trajectory, showing the path of the owl by smoothing out errors and interpolating missing detections to ensure continuity. The figure demonstrates that significant spikes in the blue trajectory are effectively filtered out, except between frames 0 and 100, where a complex background introduces substantial errors. Despite these fluctuations, the filter successfully approximates the bird’s location, even when the bounding boxes are not perfectly centered.

Another example is shown in Figure 8, featuring an eBird video (ML213887601) of a common wood pigeon (*Columba palumbus*), where significant background complexity and notable bird movements, such as picking up and eating berries, are observed.

**Figure 8.**
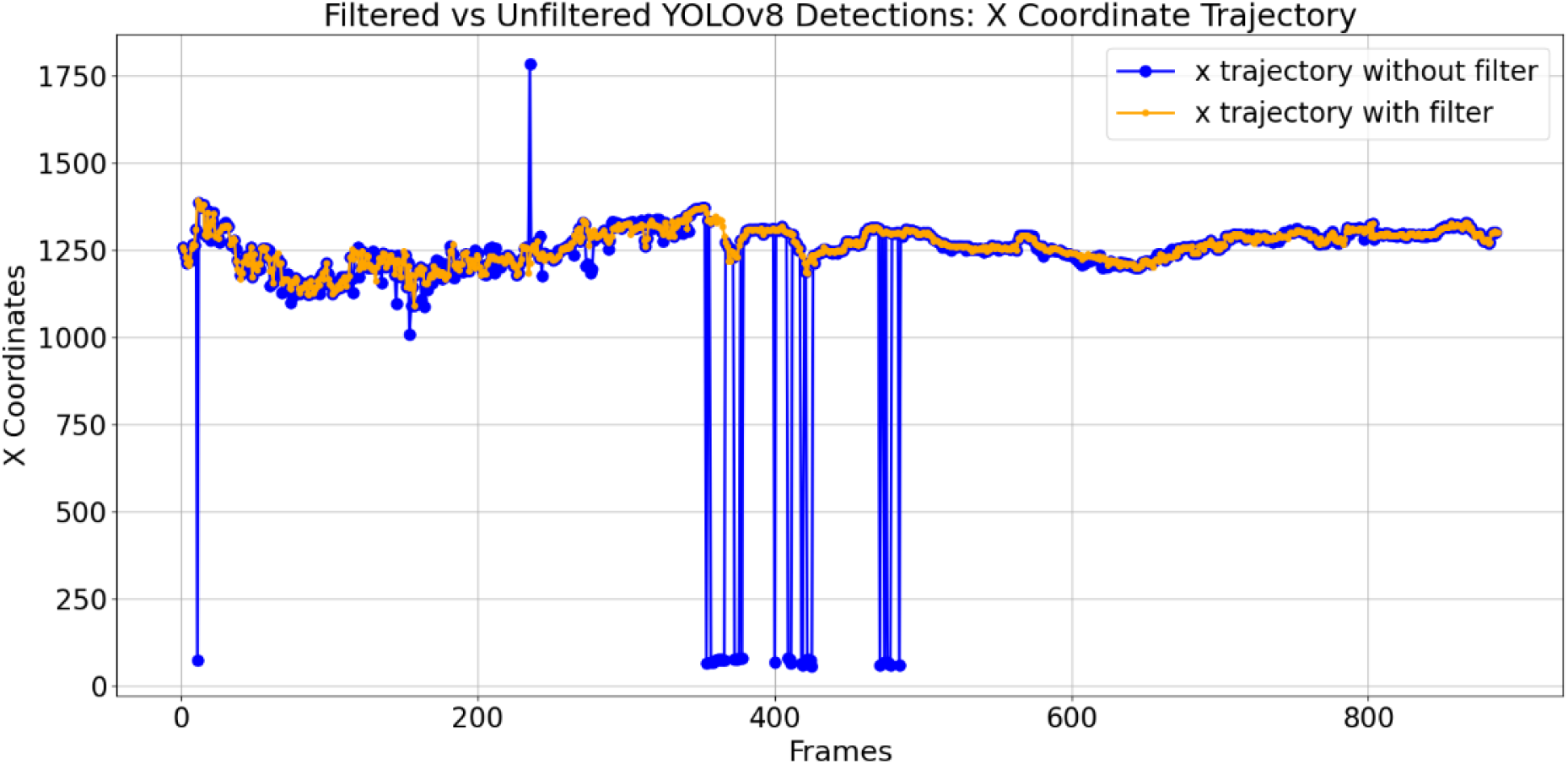
Trajectory of bounding boxes center along the X-axis across video frames for a common wood pigeon.

Figure 8 illustrates an extreme scenario with considerable incorrect bounding box estimations, particularly evident between frames 0 and 400, as indicated by the shakiness of the bounding box centers. Despite this, the filtered YOLOv8 detections successfully approximate the bird’s location. Other eBird videos with significant background complexity and unconstrained settings produced similar results to Figures 7 and 8, demonstrating the filter’s effectiveness.

### 3.2. Bird Head Pose Estimation

In this section, the results of the BHPE approach are presented, using the HRNetExp model. Sub-subsection 3.2.1 presents the results for training and validating HRNetExp on BirdGaze, testing on Birdsnap, and fine-tuning the model using the eBird video dataset. Sub-subsection 3.2.2 discusses the proposed method for refining detected bounding boxes and key points, enhancing the accuracy of head-centered detections. Sub-subsection 3.2.3 addresses the effectiveness of the key point validation step in identifying and removing errors during rapid head movements. Finally, sub-subsection 3.2.4 explores the application of key point position smoothing to minimize flickering effects and improve the analysis of head movements.

#### 3.2.1. HRNetExp Training, Validation, Testing and Fine-tuning

Figure 9 illustrates the training process of the HRNetExp model, using the BirdGaze dataset. The PCK@0.05 metric was used to evaluate the obtained accuracy, measuring the percentage of correct key points within 5% of bounding box dimensions from the ground truth.

**Figure 9.**
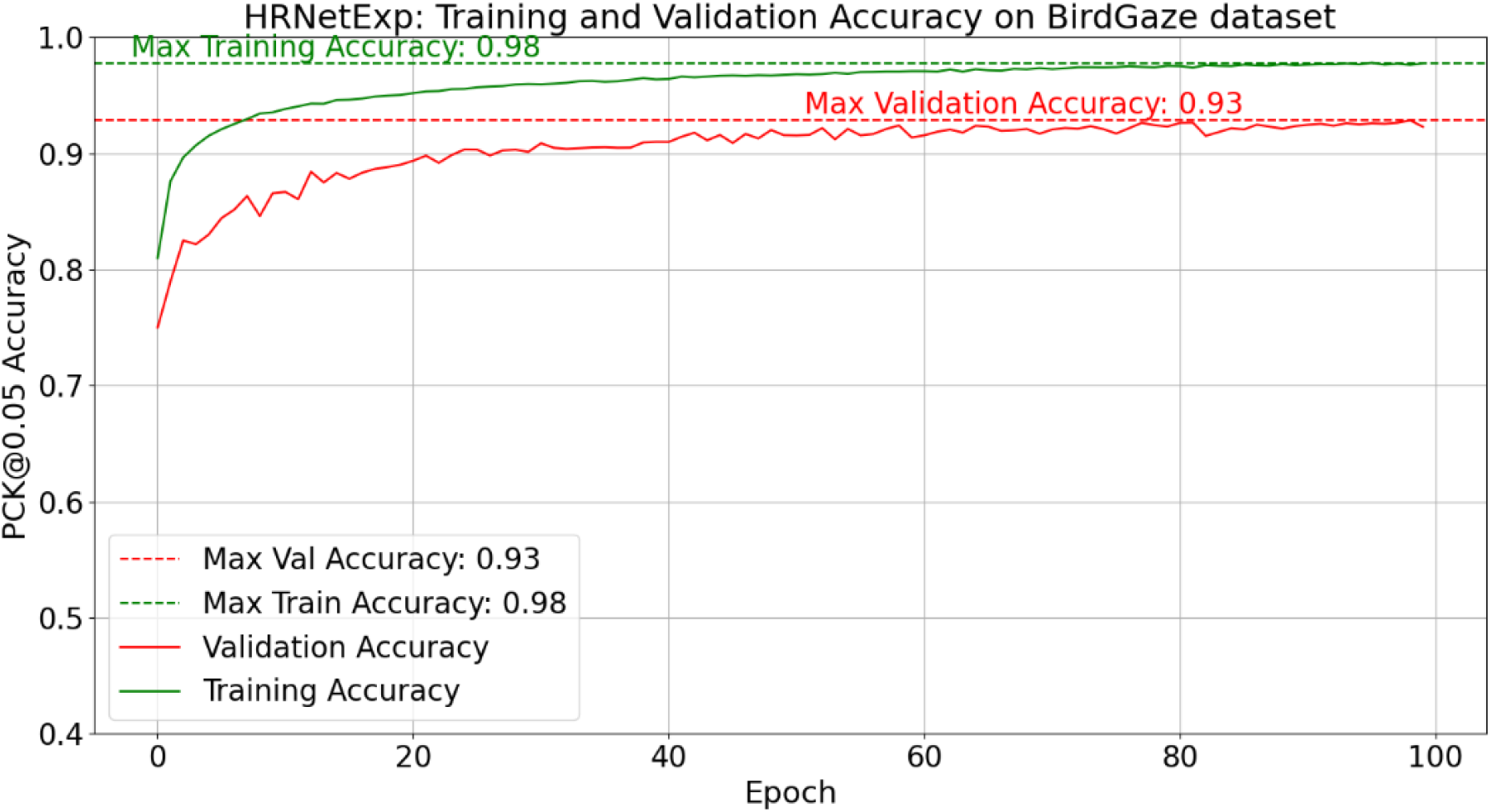
Training progress of HRNetExp on BirdGaze dataset, showcasing the evolution of training/validation accuracy across 100 epochs.

The figure showcases that the model was successfully trained without signs of overfitting on the training data. HRNetExp improved accuracy from 0.91 in HRNet4 model (trained on Animal Kingdom alone) to 0.93, indicating benefits from the larger training dataset.

The Birdsnap test subset was used to further evaluate HRNetExp, with results shown in Table 3. When analyzing Table 3, particularly the predictions with a PCK score between 0.8 and 1, reveals that HRNetExp achieves the highest number of correct key point location predictions within this range. However, there are examples with low PCK scores, showing that HRNetExp model, like was previously observed for bird detection, struggles with cases of bird occlusions and cases where birds blend into their environment.

**Table 3.** PCK scores obtained by BHPE models on Birdsnap test set.

| PCK Range | PCK > 0.8 | 0.8 ≥ PCK > 0.6 | 0.6 ≥ PCK > 0.4 | 0.4 ≥ PCK > 0 |
| --- | --- | --- | --- | --- |
| HRNet4 | 1327 (77.88%) | 294 (17.25%) | 14 (0.82%) | 69 (4.05%) |
| HRNetExp | 1420 (83.33%) | 246 (14.44%) | 10 (0.59%) | 28 (1.64%) |

After fine-tuning HRNetExp with video frames from the eBird dataset, to include examples of other bird species, the resulting model was evaluated using a separate set of 107 manually annotated eBird video frames reserved for this purpose. The results obtained before and after fine-tuning the HRNetExp model are presented in Table 4.

**Table 4.** PCK scores obtained by the HRNetExp BHPE model before and after fine-tuning, reported on the eBird test set.

| PCK Range | PCK > 0.8 | 0.8 ≥ PCK > 0.6 | 0.6 ≥ PCK > 0.4 | 0.4 ≥ PCK > 0 |
| --- | --- | --- | --- | --- |
| Not fine-tuned | 25 (23.36%) | 24 (22.43%) | 19 (17.76%) | 39 (36.45%) |
| Fine-Tuned | 95 (88.79%) | 9 (8.41%) | 3 (2.80%) | 0 (0.00%) |

The results in Table 4 show that the fine-tuned HRNetExp model performs significantly better than the non-fine-tuned version, as expected.

Future work will include manually annotating more images to better generalize across diverse bird species, poses, and challenging scenarios. Nevertheless, and whereas fine-tuning on a wider range of species can significantly improve key point identification performance, there may be unique cases for which the model may struggle with, especially given that more than 10k bird species are known and they exhibit varying behaviors.

#### 3.2.2. Bounding Box and Key Point Refining

The proposed bounding box and key point refining strategy improves estimations for birds with elongated necks, such as ostriches, where the originally detected bird bounding box sometimes excludes the bird’s head.

Figure 10 includes an example for an eBird video (ML192511121) featuring a common ostrich (*Struthio camelus*). Figure 10 compares the X-coordinate trajectories of bounding box centers provided by the bird detection module, which are body-centered, and the head-centered detections proposed in this paper.

**Figure 10.**
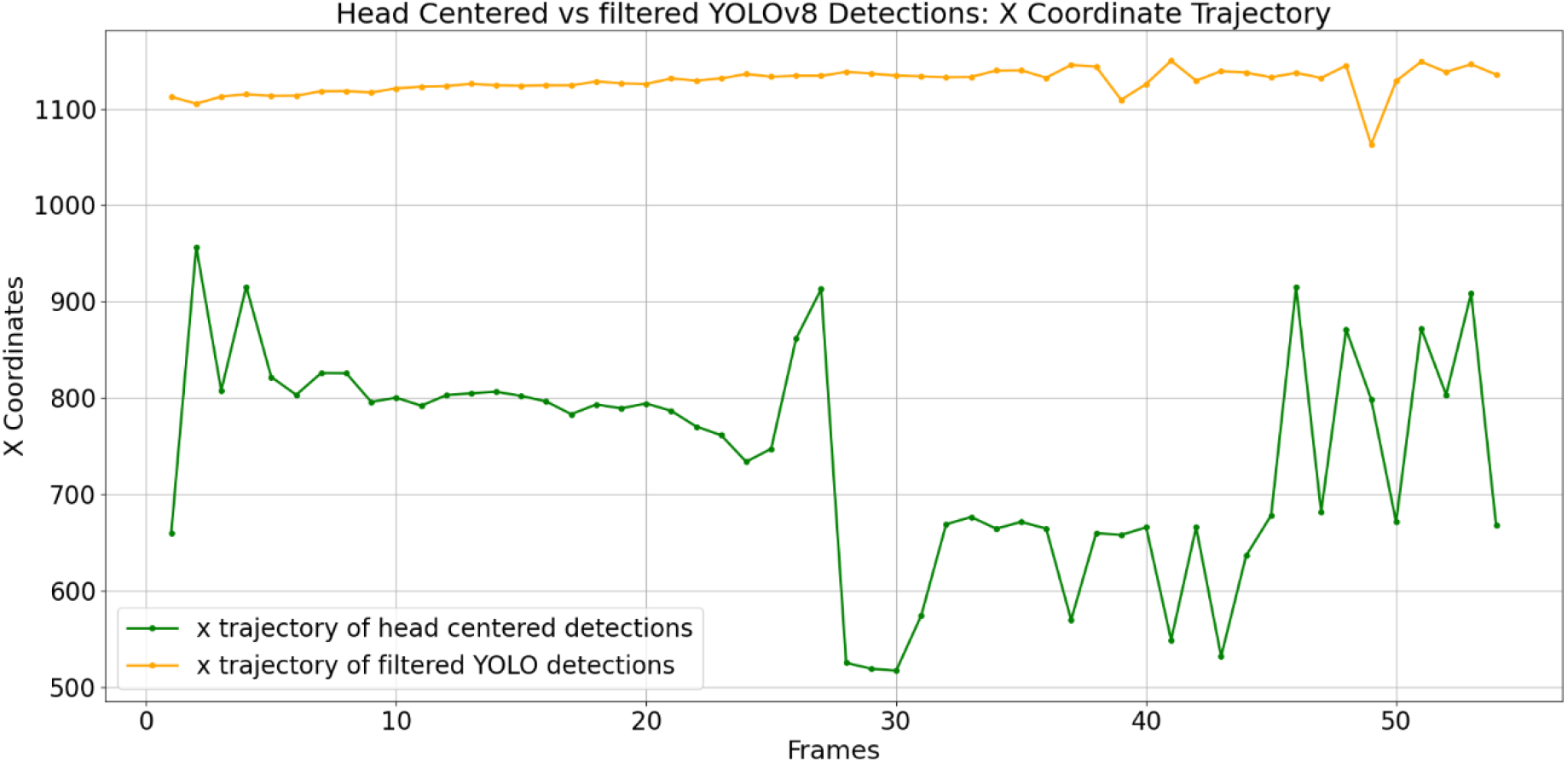
X-coordinate trajectories of bounding box centers for bird detections centered on an ostrich’s body (yellow) and head (green).

Figure 10 clearly shows a large separation between the head-centered and body-centered detections, illustrating that the YOLOv8-based bird detection model adopted often excluded the ostrich’s head from the detected bounding box. However, the head-centered detections are not perfectly aligned with the ostrich’s head, evidenced by the spikes in the green trajectory, due to initial key point errors occurring in frames where the head is initially excluded. Despite these errors, the head-centered detections successfully capture the ostrich’s head, improving the model’s effectiveness in such cases.

To illustrate the behavior of the proposed trajectory refinement procedure, Figure 11 compares the estimated head and beak key point locations, with and without refining. The figure shows a first step of key point estimation on frames centered on the ostrich’s body (green), and a refined key point estimation obtained from frames centered on the ostrich’s head (yellow). The unrefined green line shows deviations caused by head exclusion, while the refined yellow line demonstrates smoother trajectories. Remaining spikes are due to unseen pose-related errors

**Figure 11.**
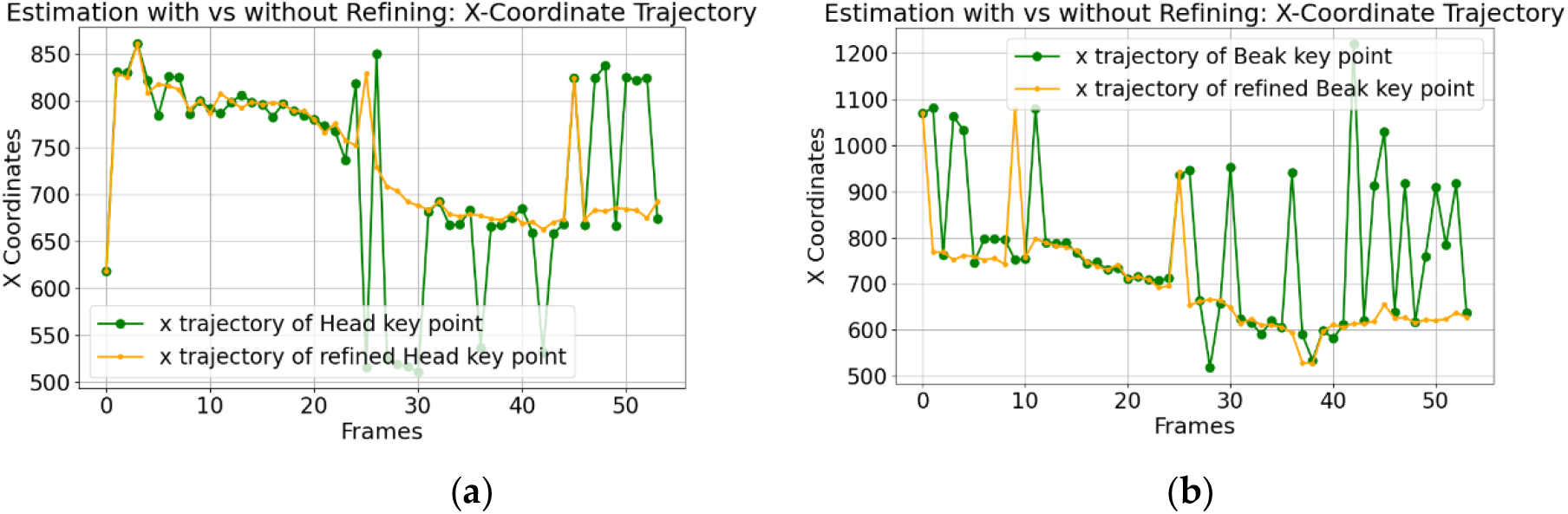
X-coordinate trajectory of the ostrich estimated key points.: (**a**) Head key point.; (**b**) Beak key point.

Despite improvements, some pose-related errors may remain, as illustrated in Figure 12, because certain poses were not well represented in the training set, highlighting the need for more training data.

**Figure 12.**
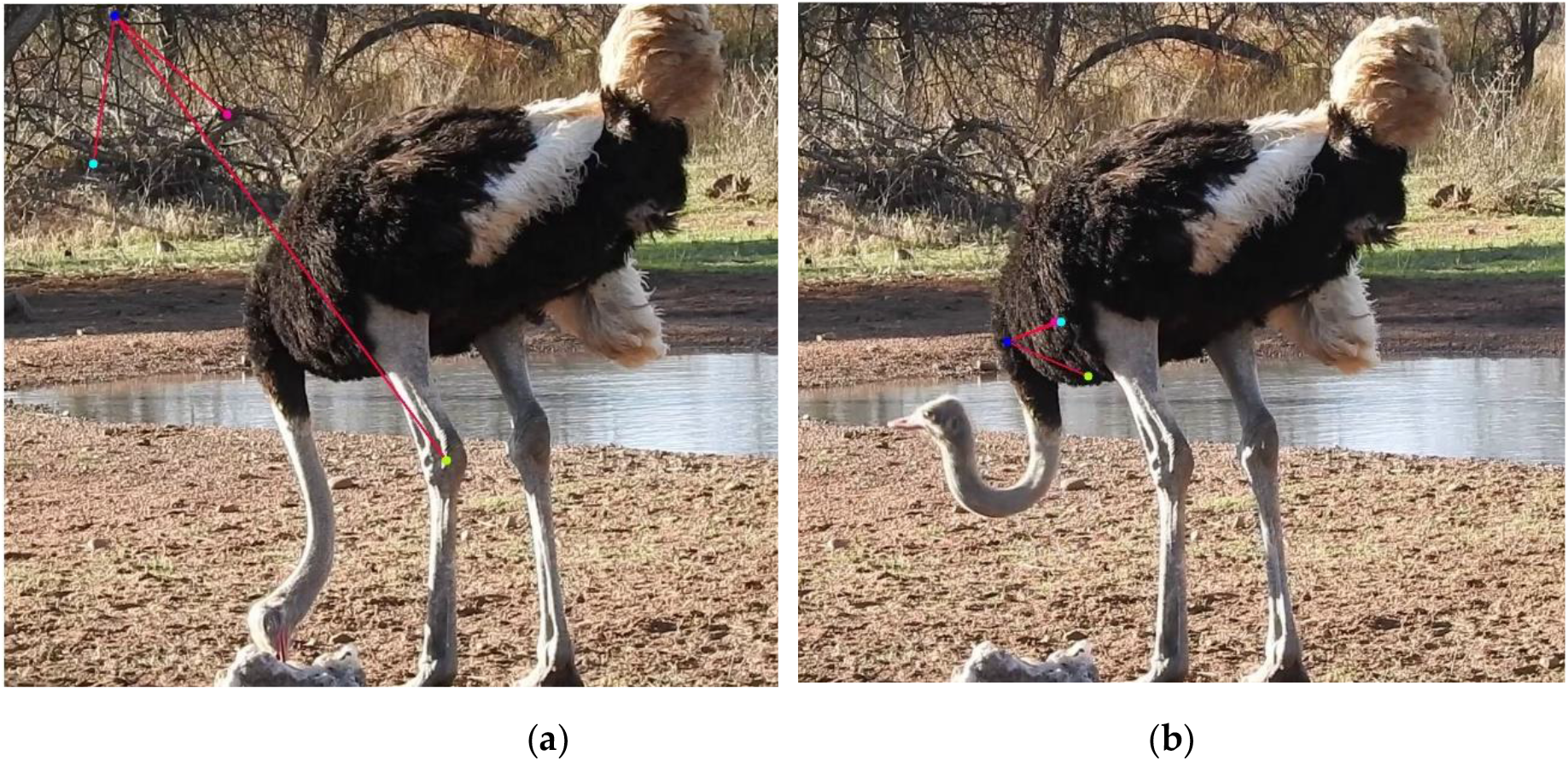
Persistent pose-related errors of head pose estimation, using key point refinement.: (**a**) Beak occlusion.; (**b**) Leaning posture.

Figure 13 shows that, on average, key point estimation accuracy significantly improves with the proposed refinement procedure. This approach effectively corrects most head exclusions and leads to higher average HRNetExp confidence scores for each estimated key point.

**Figure 13.**
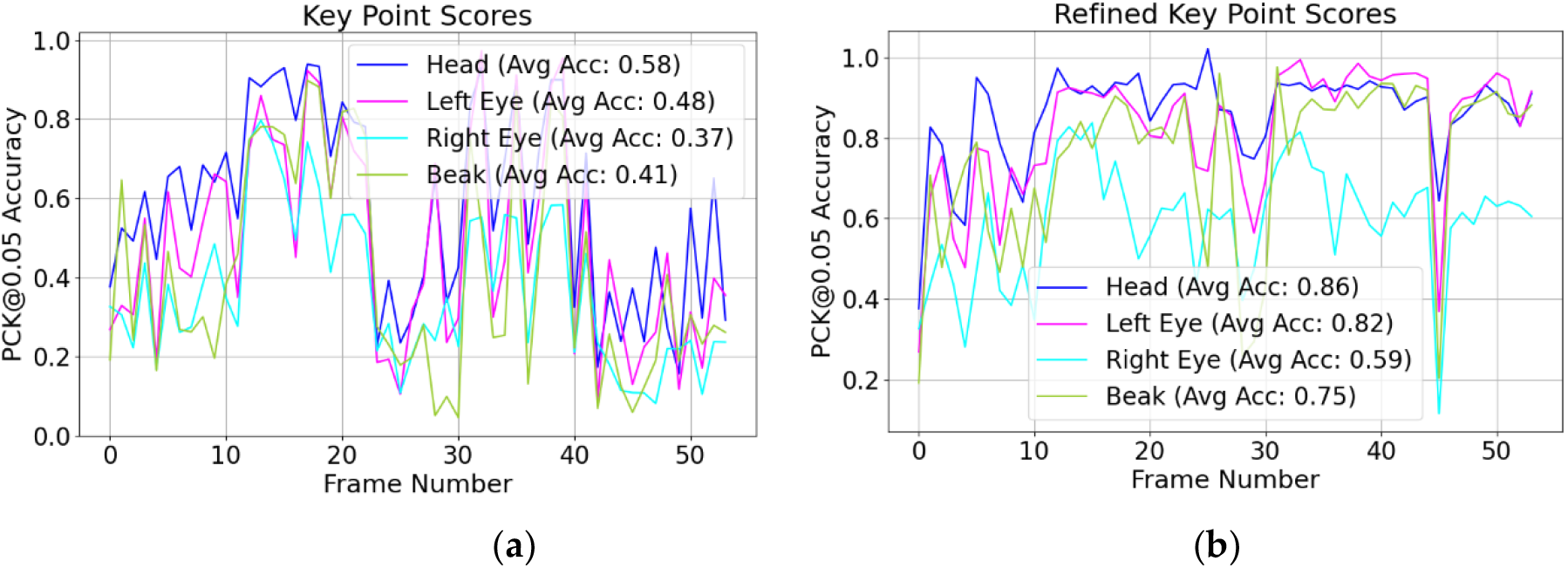
Key point estimation scores throughout the video of a common ostrich.: (**a**) Scores without refining.; (**b**) Scores with refining.

In conclusion, testing across various videos from the eBird dataset consistently showed improvements in accuracy of estimations. Therefore, it can be concluded that incorporating the key point refining step offers substantial benefits compared to not using it.

#### 3.2.3. Key Point Validation

To illustrate the usefulness of the proposed key point validation step, an example of an eBird video of a barn owl (*Tyto alba*) (ML194115401) performing rapid head rotations was analyzed. Figure 14 shows the applied distance and score thresholds.

**Figure 14.**
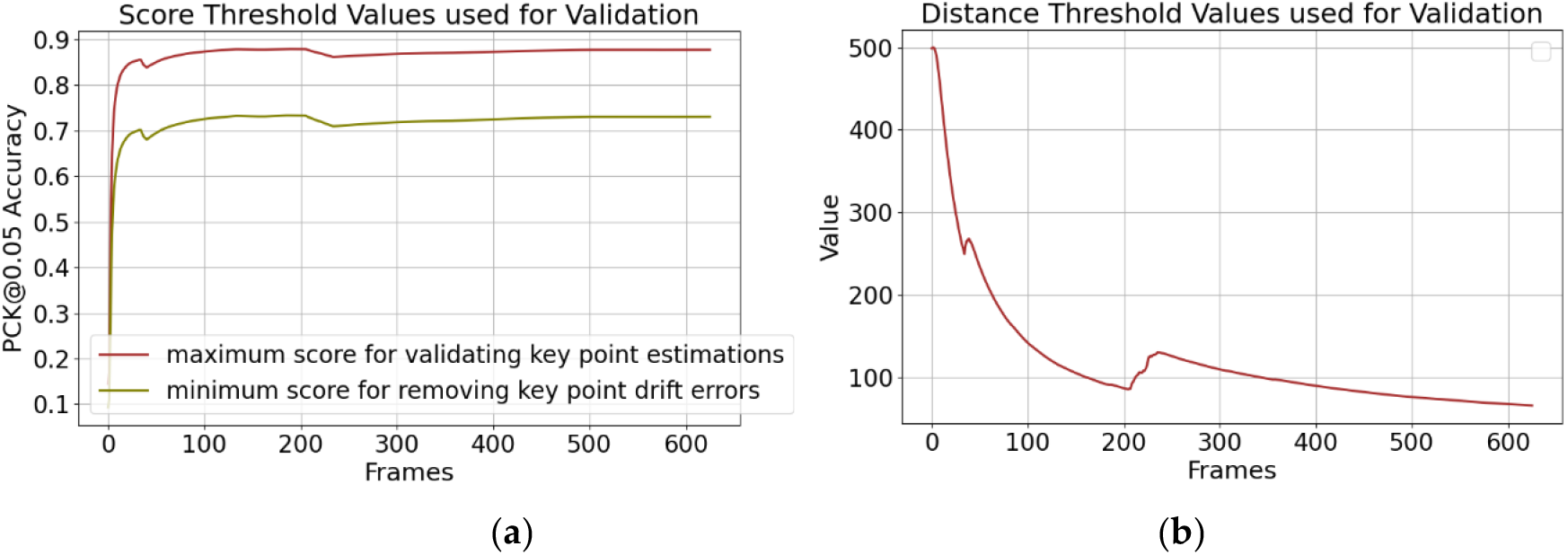
Threshold values for key point validation.: (**a**) Maximum and minimum score thresholds.; (**b**) Absolute distance threshold.

Figure 14 indicates that the score thresholds start at lower values due to initial key point estimation errors but later stabilize at 0.73 for the minimum and 0.87 for the maximum thresholds. Likewise, the distance threshold begins high at 500 because of early errors, decreases slightly, then rises again with subsequent errors, ultimately stabilizing as these errors diminish.

To demonstrate the effectiveness of error detection and removal, Figure 15 presents the X-coordinate trajectory of the owl’s head and beak key points.

**Figure 15.**
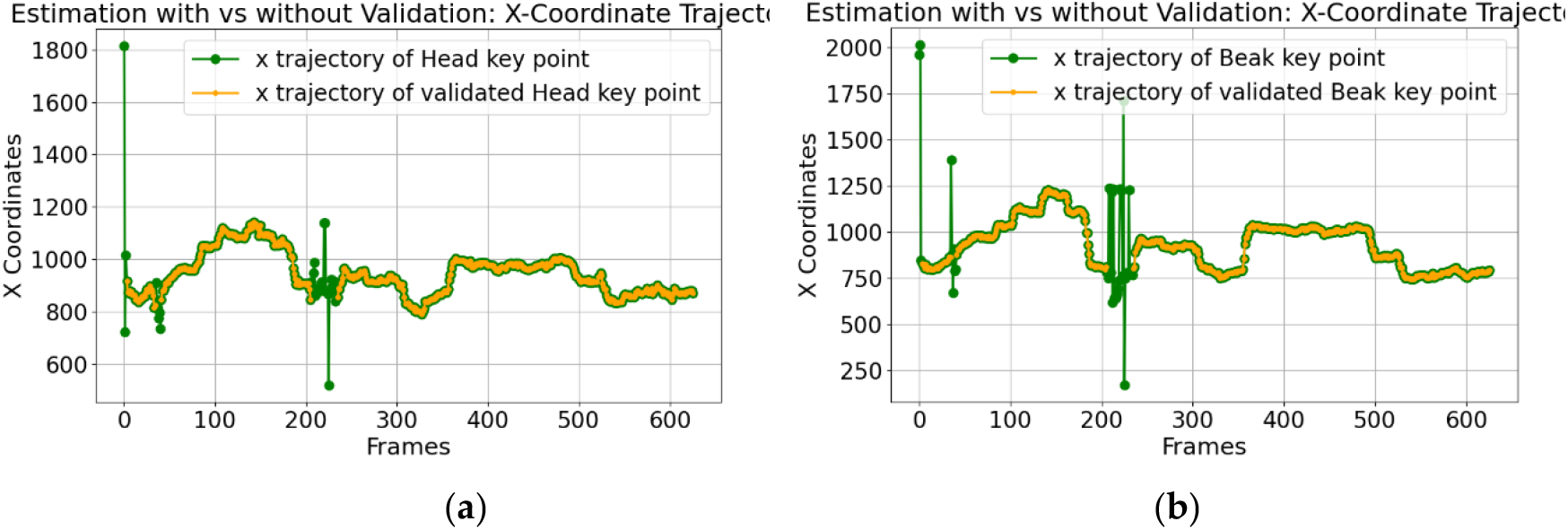
X-coordinate trajectory of the barn owl estimated key points.: (**a**) Head key point; (**b**) Beak key point.

The figure highlights significant aberrant spikes in the key point estimations that signify estimation errors (green). These spikes are identified as errors and then removed (yellow). Closer examination of errors around frame 0 and frame 225 reveals they stem from rapid head movements, resulting in motion blur due to the video acquisition frame rate (see Figure 16). The owl’s quick movements appear as blurred streaks, making it difficult for the model to accurately detect and estimate key points on the bird’s head.

**Figure 16.**
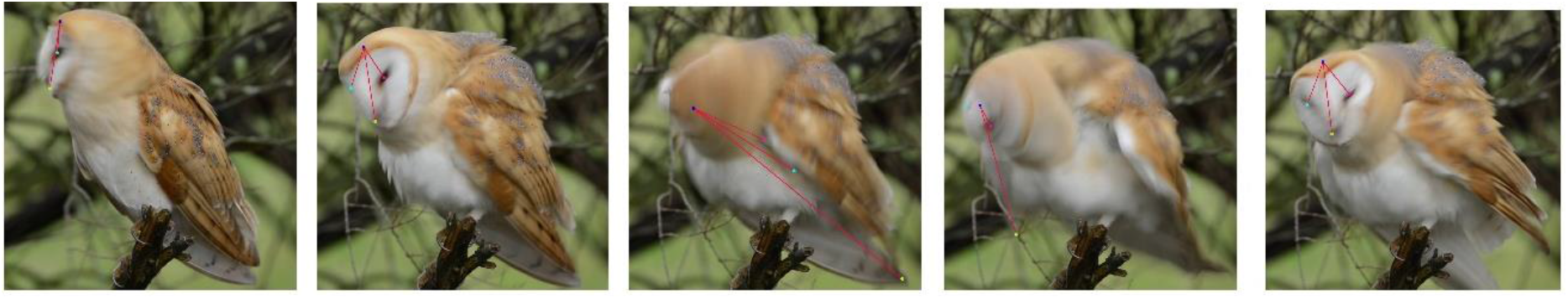
Motion blur caused by low video frame rate, during rapid head movements of the owl, showing errors in key point estimation.

In conclusion, the video frame rate causes motion blur during rapid head movements, leading to errors in key point estimations. Whereas these errors are effectively detected and removed by the filter, a higher frame-rate video would have enabled correct estimations during fast head rotations.

#### 3.2.4. Key Point Position Smoothing

An eBird video (ML608894819) of a walking common wood pigeon (*Columba palumbus*), performing subtle head movements, was analyzed to evaluate the Key Point Smoothing step. Figure 17 shows the computed head angular displacement with and without smoothing. The smoothing filter appears to reduce most spikes likely caused by flickering, while retaining general movement patterns, though this cannot be fully confirmed without ground truth data. The remaining spikes, which did not meet the smoothing criteria are considered true head movements.

**Figure 17.**
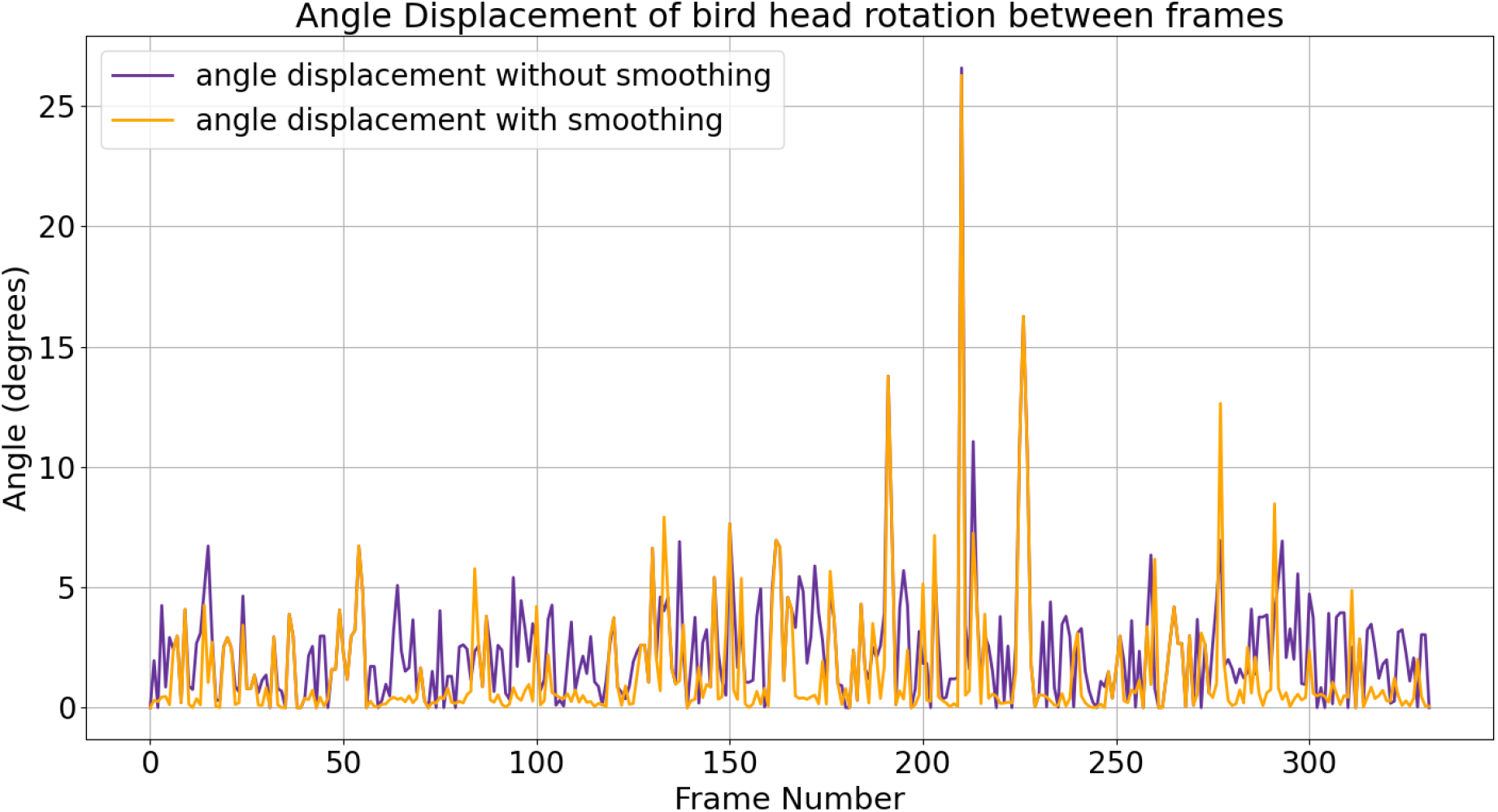
Angle displacement of bird head rotations across consecutive frames, measured between head and beak key points. The figure shows noise reduction (orange line) compared to the original noisy measurements (purple line).

Figure 18 shows the frequency spectrum of bird head motion with and without smoothing. In Figure 18(a), flickering introduces aberrant spikes at higher frequencies, where flickering noise concentrates. Applying the smoothing filter reduces noise by lowering amplitudes across frequencies, but while most noise is reduced, complete elimination could risk filtering out genuine head movements.

**Figure 18.**
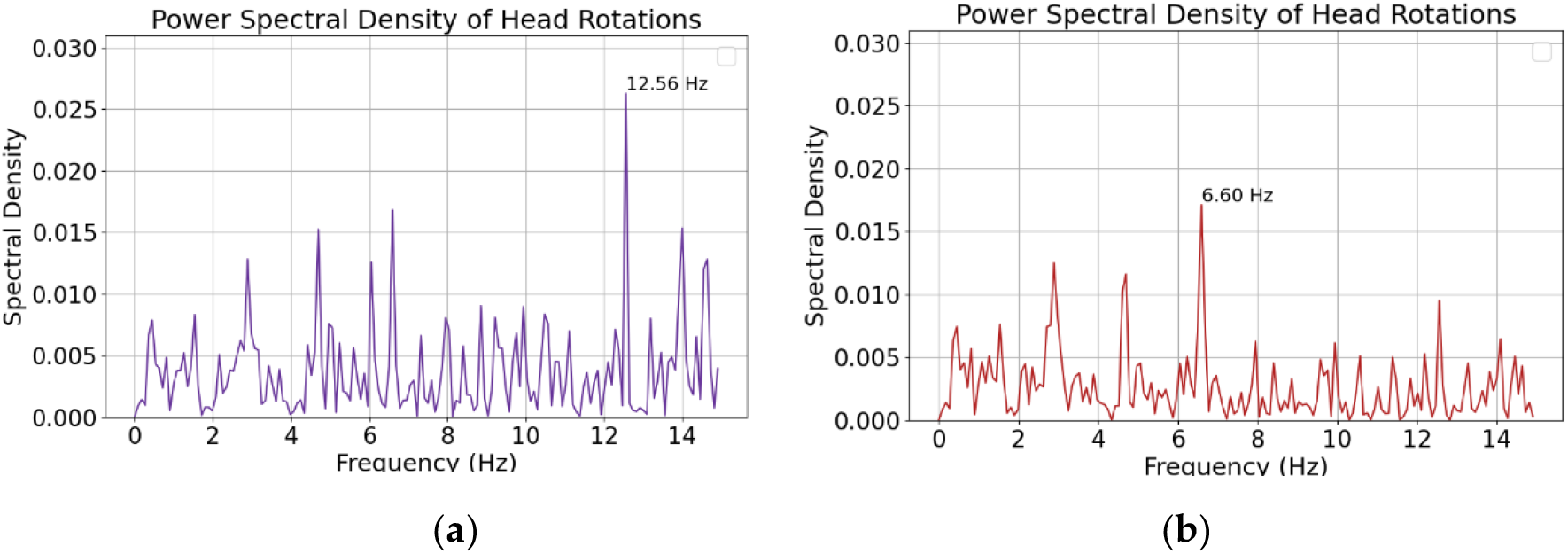
Comparison of power spectral density with and without smoothing.: (**a**) Without smoothing; (**b**) With smoothing.

In conclusion, while key point smoothing does not eliminate noise caused by flickering, it significantly reduces it to a negligible level, allowing the predominant head motion to be preserved.

### 3.3. Head Motion Computation

In this section, results of head motion computations are presented. Sub-subsection 3.3.1 presents results of the analysis of bird head motion on eBird videos, using 2D key points obtained through the proposed methodology, and addresses challenges posed by video frame rates, camera angles and noise. Sub-section 3.3.2 presents results of analyses on the 3D-POP dataset, again using 2D key points obtained through the proposed methodology. This dataset provides 2D and 3D annotated videos of pigeons, enabling direct comparison between estimated head motion and ground truth, which serves to assess the accuracy of the proposed methodology in the 2D domain. Sub-subsection 3.3.3 discusses the transition to the 3D domain using the 3D-POP dataset.

#### 3.3.1. 2D Realm - eBird videos

An eBird video (ML201912681) of a common buzzard (*Buteo buteo*) was analyzed to compute head motion, specifically focusing on the frequency of head rotations in the pitch direction. The video, recorded at 30 fps, shows the bird performing moderate head rotations while scanning for prey. The computed PSD of these head rotations is shown in Figure 19. While a distinct peak at 0.31 Hz is observed, it is important to acknowledge that these results merely demonstrate the framework’s ability to produce outputs, not necessarily validated head motion. The absence of peaks around 14 Hz suggests the flickering issue has been mitigated, but it cannot be definitively concluded that all noise has been eliminated. The remaining peaks may reflect either actual bird motion or residual noise.

**Figure 19.**
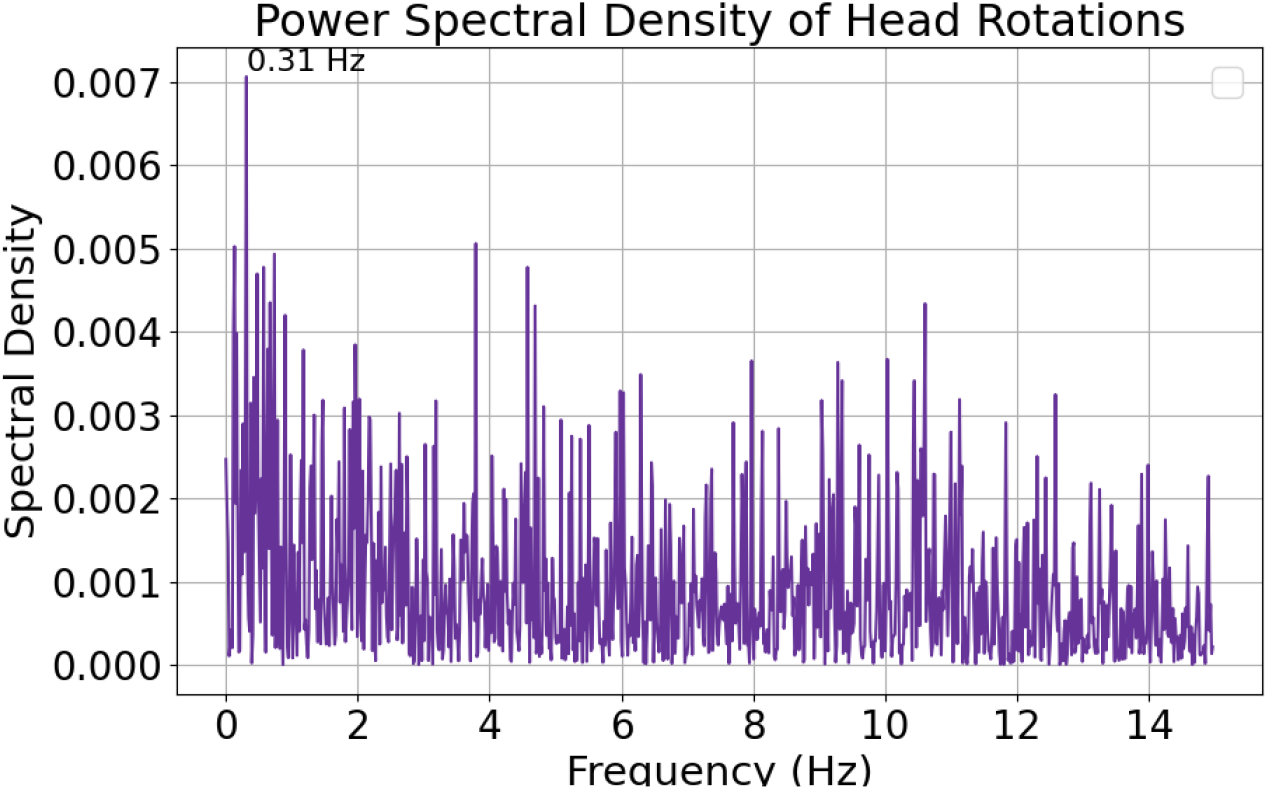
Power spectral density of head rotations from a common buzzard performing a moderate-speed head rotation. The figure shows a distinct frequency peak, around 0.31 Hz.

To further explore this, another eBird video (ML201465631) of a great crested grebe (*Podiceps cristatus*) performing subtle head rotations was analyzed (Figure 20). This video, recorded at 25 fps, shows that the smoothing filter reduced high-frequency noise, as evidenced by the absence of high-amplitude peaks. However, the observed peak frequency at 0.27 Hz remains in a range less affected by noise.

**Figure 20.**
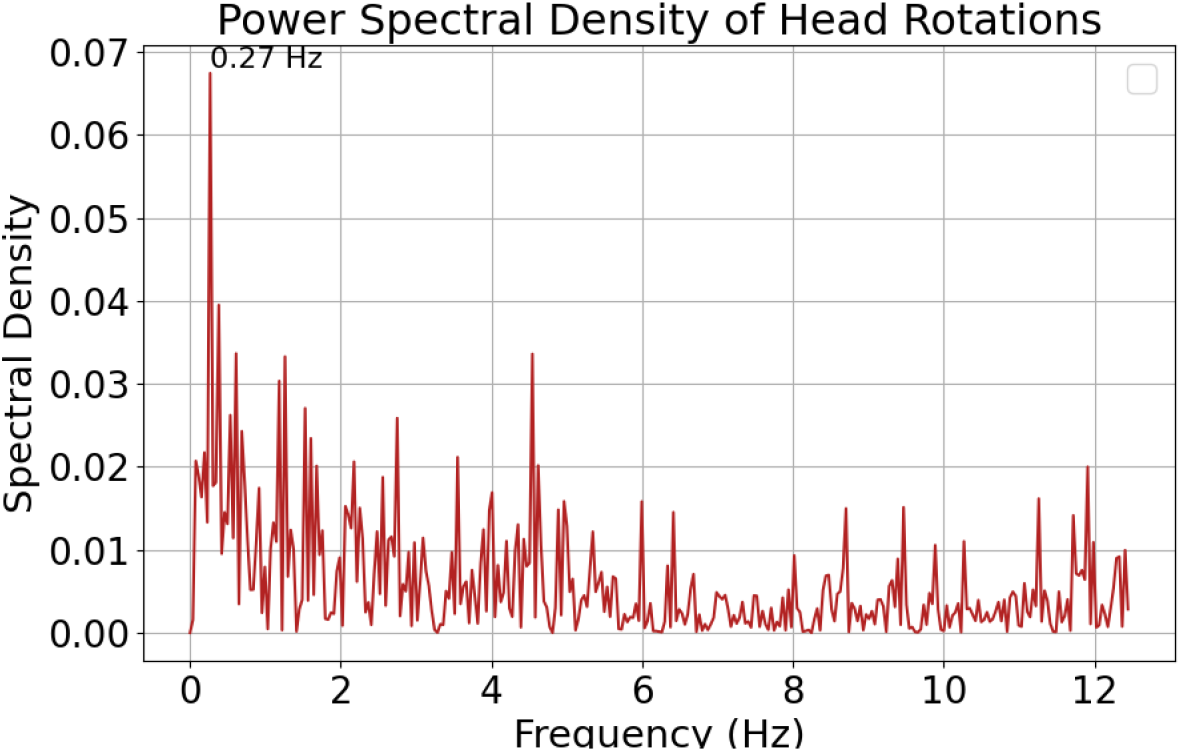
Power spectral density of head rotations from a great crested grebe (*Podiceps cristatus*) performing subtle head rotations. The figure shows a distinct frequency peak, around 0.27 Hz.

This part of the analysis demonstrates that the framework can process head motion data and produce plausible outputs, but without ground truth validation, these results must be interpreted with caution. The limitations of 2D representation, video frame rate, bird pose relative to the camera, and noise from the 2D BHPE model significantly impact the accuracy of the findings. While these results indicate the feasibility of head motion analysis, they primarily serve as a precursor to more robust evaluations, such as comparisons with actual validated head motion data, which are explored in the following sections.

#### 3.3.2. 2D Realm – 3D POP videos

To validate the proposed methodology in the 2D domain, we first estimated key points on one randomly selected video from the 3D-POP video dataset. We then computed head motion metrics from these key points and compared them to head motion metrics computed from 2D manual annotations (control). Our analysis reveals that both estimated and control head motion showcase a maximum frequency peak at 2 Hz, with a very similar low frequency pattern (Fig. 21). However, compared to control head motion, the estimated one shows higher peaks in the higher frequencies (> 8Hz), indicating that, though minimized, noise was not entirely removed by the proposed filtering methodology.

**Figure 21.**
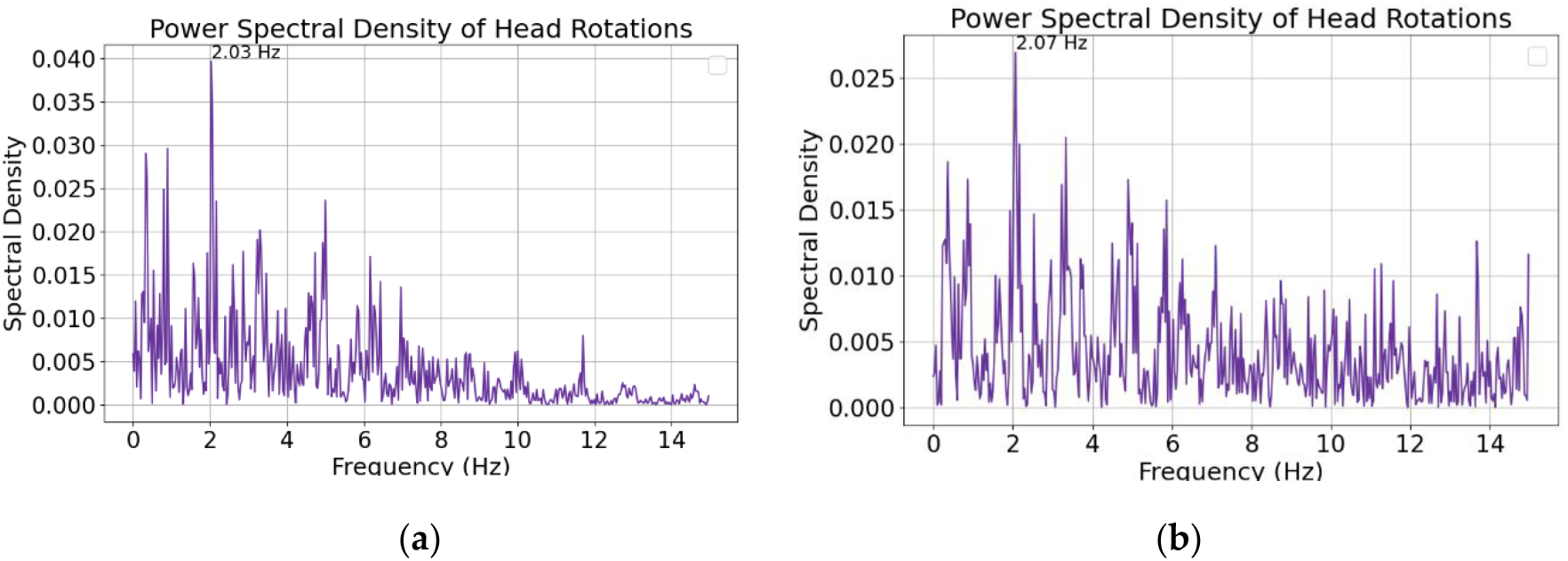
Comparison of power spectral density obtained with control and estimated data.: (**a**) Control; (**b**) Estimated.

#### 3.3.3. 3D Realm - 3D POP videos

To validate and extend the proposed methodology for head motion analysis to the 3D realm, we analyzed one 3D video set, randomly selected from the 3D-POP dataset, featuring a common wood pigeon (*Columba palumbus*) walking and feeding on the ground. The analysis focuses specifically on head motion and reveals consistent rotation patterns in pitch, roll, and yaw (Fig. 22). Head motion is observed to be maximal in the low frequencies and reduced in the high frequencies when compared to analyses restricted to the 2D realm (Fig. 23). It is important to clarify that this analysis pertains solely to head motion and does not involve testing the AI model in 3D. These results highlight that our head rotation estimation methodology can successfully be applied to the 3D realm, with clear noise reduction, resulting in more robust results than in the 2D realm.

**Figure 22.**
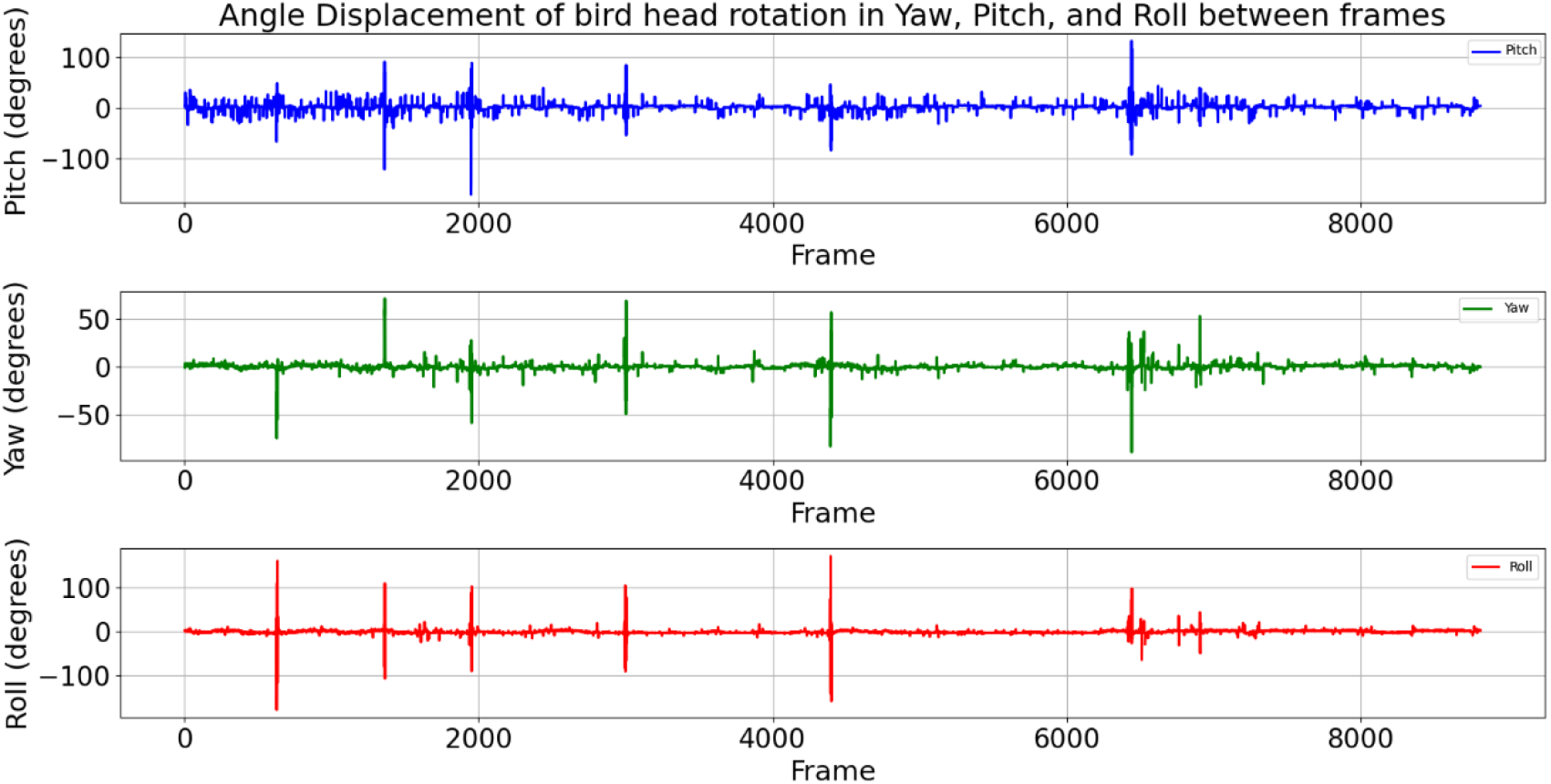
Angular displacement of a bird head, computed from 3D annotated frames from the 3D-POP dataset along the head-centered pitch (blue), roll (red) and yaw (green) axes rotation.

**Figure 23.**
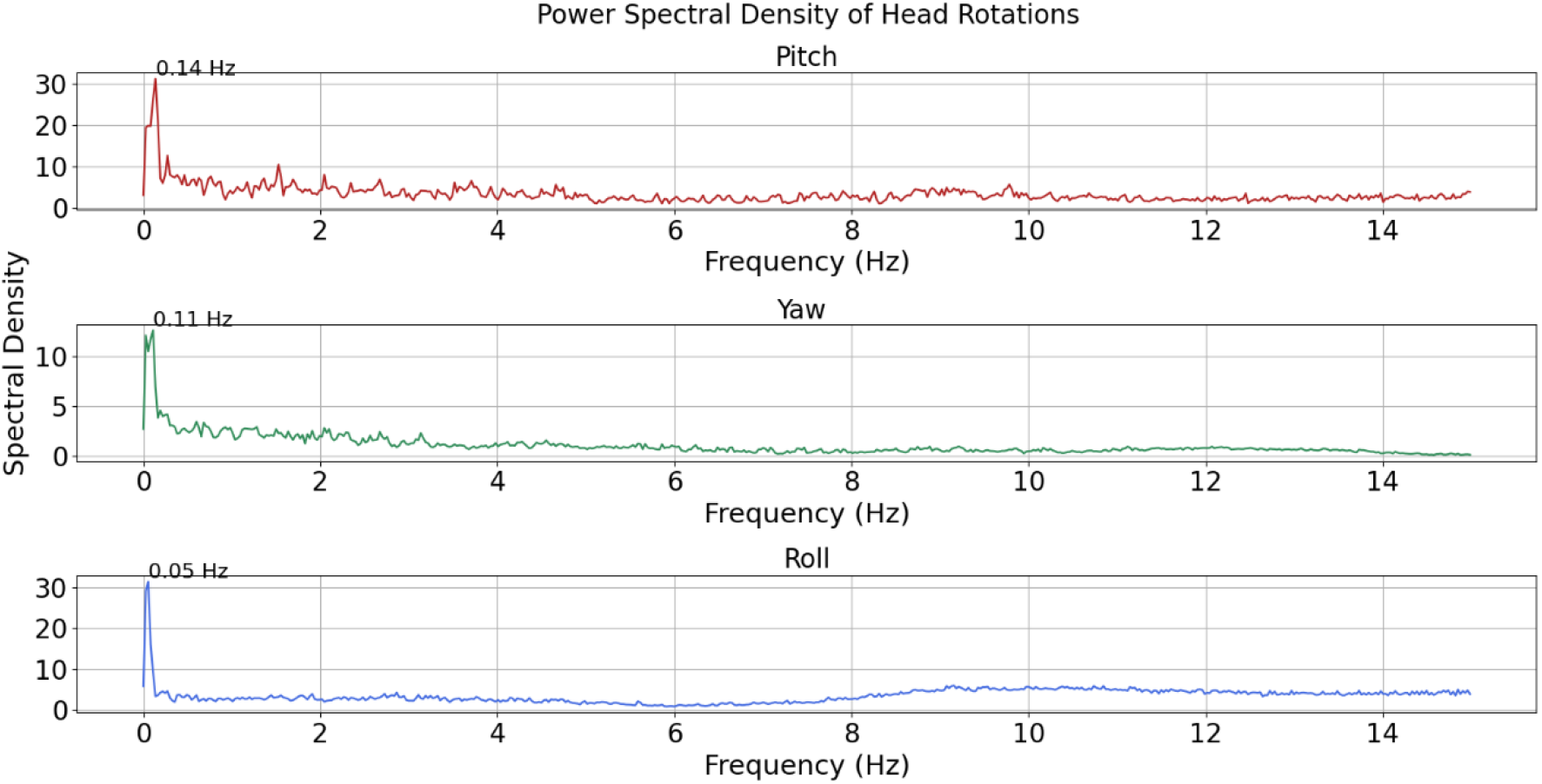
Power spectral density of rotations of a bird head, computed from 3D annotated frames from the 3D-POP dataset along the head-centered pitch (red), roll (blue) and yaw (green) axes rotation.

## 4. Discussion

This study presents a new framework for analyzing bird head motion from videos, using a 2D BHPE model trained on the BirdGaze dataset, with potential adaptability to the 3D realm. It demonstrates that bird head motion can be analyzed, though at this point with some limitations. These include the quality and quantity of available high-frame rate bird video data, as the lack of sufficient annotated videos makes it challenging to compare head motion estimates to ground truth. These limitations highlight the need for more comprehensive and higher-quality datasets to generalize and improve the accuracy of the proposed approach.

Current datasets used to train BHPE models primarily focus on 2D key point estimation, limiting their ability to capture the full spectrum of 3D rotations, including yaw, pitch, and roll. This reliance on 2D key point estimation confines the analysis to videos in which the bird faces the camera from the side. While this study selected such videos through visual inspection, future efforts could automate this process by enhancing 2D datasets.

The inclusion of bird videos from platforms like eBird offers a rich resource, but these videos are often captured in uncontrolled environments, where complex backgrounds and lighting conditions can introduce errors in key point estimation. Furthermore, the critical goal of this research is to analyze the frequencies and angular velocities of volitional bird head rotations, yet the high-speed, quick-turning behavior of many bird species—of which there are over 10,000—remains underexplored due to frame rate limitations. Most available videos are recorded at standard frame rates, which limit the range of detectable frequencies and reduce the accuracy of head motion analysis. Using high-speed cameras to record bird movements at higher frame rates would provide the granularity needed to accurately measure rapid angular velocities and detect higher frequency head motions. This would allow for a more reliable analysis of fast bird head movements. Additionally, a key focus should be on head motion occurring during locomotion, as locomotion plays a central role in the adaptation of semicircular ducts. While this analysis includes various behaviors, prioritizing locomotion-related head movements would offer deeper insights into the relationship between head motion dynamics and vestibular system adaptations.

Another limitation in this study is the reliance on a 2D BHPE model for head motion analysis, which restricts estimations to a single view. While the model demonstrated high accuracy in estimating key points in individual frames, it introduced a flickering effect between consecutive video frames, which added noise to the head motion calculations. Despite efforts to eliminate this noise using a smoothing filter, it could only be minimized, not fully removed. Applying a stronger filter would completely eliminate the noise, but at risk of also removing actual head movements. As a result, the minimized noise is still visible in the frequency spectrum, limiting the range of frequencies that can be accurately analyzed without interference. Due to the Nyquist limit, only half of the recorded frame rate can be analyzed, and with noise affecting approximately half of those frequencies, only about one-quarter of the total frame rate is effectively usable. For instance, to accurately analyze head motion up to 20 Hz, a frame rate of at least 80 Hz would be necessary to ensure sufficient signal clarity and frequency resolution. Future research should prioritize acquiring higher-quality datasets, particularly videos recorded at higher frame rates, to improve the accuracy of head motion analysis and ensure reliable detection of rapid angular velocities and higher frequency motions. Additionally, further exploration of smoothing methods, such as applying filters at the level of the power spectral density (PSD) rather than directly on the key points, could help reduce noise more effectively while preserving true motion signals.

The analysis highlights the significant impact of noise in 2D key point positions and underscores the importance of high video frame rates to accurately detect strenuous head motion. Noise can greatly hinder the accurate analysis of head motion frequencies by introducing spurious frequency peaks that may resemble actual head movements. Additionally, the frame rate limitations restrict the ability to capture higher frequencies, further constraining the analysis.

## 5. Conclusions

This study presented a framework for bird head motion analysis using a 2D BHPE model trained on the BirdGaze dataset. The framework proved effective in estimating key points in 2D videos. The approach also demonstrated potential for 3D applications, although currently limited to head motion calculations based on manually annotated key points. Certain limitations, such as reliance on 2D data, noise and challenges posed by uncontrolled recording environments, were identified. Despite these constraints, this framework enables analyzing bird head rotations and corresponding frequencies, contributing valuable insights. Addressing limitations by integrating 3D key point estimation and high-speed recordings could further refine the analysis and expand its applicability. The study lays the groundwork for future advancements in bird head motion studies, with the potential to enhance understanding of avian dynamics.

